# Prolonged β-Adrenergic Stimulation Disperses Ryanodine Receptor Clusters in Cardiomyocytes: Implications for Heart Failure

**DOI:** 10.1101/2022.02.18.481024

**Authors:** Xin Shen, Jonas van den Brink, Anna Bergan-Dahl, Terje R. Kolstad, Einar S. Norden, Yufeng Hou, Martin Laasmaa, Ann P. Quick, Emil K.S. Espe, Ivar Sjaastad, Xander H.T. Wehrens, Andrew G. Edwards, Christian Soeller, William E. Louch

## Abstract

Ryanodine Receptors (RyRs) exhibit dynamic arrangements in cardiomyocytes, and we previously showed that “dispersion” of RyR clusters disrupts Ca^2+^ homeostasis during heart failure (HF) (Kolstad *et al*., eLife, 2018). Here, we investigated whether prolonged β-adrenergic stimulation, a hallmark of HF, promotes RyR cluster dispersion, and examined the underlying mechanisms. We observed that treatment of healthy rat cardiomyocytes with isoproterenol for 1 hour triggered progressive fragmentation of RyR clusters. Pharmacological inhibition of CaMKII reversed these effects, while cluster dispersion was reproduced by specific activation of CaMKII, and in mice with constitutively active Ser2814-RyR. A similar role of protein kinase A (PKA) in promoting RyR cluster fragmentation was established by employing PKA activation or inhibition. Progressive cluster dispersion was linked to declining Ca^2+^ spark fidelity and magnitude, and slowed release kinetics from Ca^2+^ propagation between more numerous RyR clusters. In healthy cells, this served to dampen the stimulatory actions of β-adrenergic stimulation over the longer term, and protect against pro-arrhythmic Ca^2+^ waves. However, during HF, RyR dispersion was linked to impaired Ca^2+^ release. Thus, RyR localization and function are intimately linked via channel phosphorylation by both CaMKII and PKA which, while finely tuned in healthy cardiomyocytes, underlies impaired cardiac function during pathology.

**Significance statement:** The heartbeat is triggered by the release of Ca^2+^ from Ryanodine Receptors (RyRs) within cardiomyocytes. Recent data indicate RyR arrangement is highly malleable. However, mechanisms controlling RyR reorganisation and the subsequent impact on Ca^2+^ homeostasis remain unclear. Here, we show that prolonged β-adrenergic stimulation causes RyR clusters to disperse, drastically altering the frequency and kinetics of Ca^2+^ release events called “Ca^2+^ sparks” in a process that is dependent on CaMKII and PKA. In healthy cells, these compensatory effects protect against arrhythmogenic Ca^2+^ over-activity. However, during heart failure, RyR hyper-phosphorylation and dispersion impairs Ca^2+^ release and cardiac performance. Thus, RyR localization and function are intimately linked via channel phosphorylation which, while finely tuned in health, underlies impaired cardiac function during pathology.

## Introduction

In cardiomyocytes, the processes underlying initiation of contraction are well described, at least at the macroscale. Sarcolemmal depolarisation triggers an influx of Ca^2+^ through voltage-gated L-type Ca^2+^ channels (LTCCs) within t-tubules, which in turn elicits a much larger release of Ca^2+^ into the cytosol via ryanodine receptors (RyRs) in the sarcoplasmic reticulum (SR). This process of Ca^2+^-induced Ca^2+^ release (CICR) depends on the precise organization of LTCCs and RyRs within dyadic junctions of the two membranes, with LTCCs located in nanoscale apposition to RyRs across the dyadic cleft^1^. Intrinsically, RyRs are organized into discrete clusters, and these arrangements are thought to be critical in defining channel function. Indeed, RyR clusters located within close proximity are proposed to cooperatively operate as Ca^2+^ Release Units (CRUs) to generate Ca^2+^ sparks^2,3^, the elementary units of SR Ca^2+^ release^4^. Recent advances in high-resolution imaging techniques such as 3D dSTORM^5^, DNA-PAINT^6^ and electron tomography^7^ have created an opportunity to understand these dyadic arrangements in unprecedented detail.

Contractile dysfunction and arrhythmogenesis are hallmarks of heart failure (HF), and considerable data have linked these phenomena to t-tubule disruption in this condition^8-10^. Recently, we reported that pathological changes occur also on the other side of the dyad, as we observed “dispersion” of RyR clusters in failing cardiomyocytes. This RyR rearrangement specifically included fragmentation of RyR groupings into more numerous, smaller clusters, without any change in overall channel number^11^. RyR dispersion was critically linked to low fidelity spark generation. Furthermore, when Ca^2+^ release was successfully triggered by a CRU, propagation of Ca^2+^ between multiple clusters generated sparks with slow kinetics, and the overall Ca^2+^ transient was desychronized^11^. Similar fragmentation of CRUs has been reported in a sheep model of persistent atrial fibrillation^12^, and in monocrotaline-induced right ventricular failure in rats^13^. Thus, accumulating evidence indicates that RyR mislocalization, and ensuing dysfunction, is a key contributor to pathophysiology.

What drives changes in RyR organization in diseased cardiomyocytes? Previous work has indicated that nanoscale RyR positioning is fine-tuned by phosphorylation of the channels, at least acutely^7,14^, and that both CaMKII- and PKA-dependent phosphorylation of RyR disrupt its function during HF^15-17^. We presently hypothesized that these two phenomena are intimately linked. We demonstrate that prolonged β-adrenergic stimulation, as is well known to occur in HF, promotes RyR dispersion via both CaMKII- and PKA-dependent phosphorylation of the channel. This RyR reorganization and sensitization is associated with a time-dependent increase in Ca^2+^ leak, slowing of Ca^2+^ spark kinetics, and reduced Ca^2+^ transient magnitude. In healthy cells, these actions appear to be aimed at gradually countering the stimulatory effects of increased RyR phosphorylation and Ca^2+^ sensitivity. Indeed, we observed that sufficiently dispersed RyRs inhibit the development of Ca^2+^ waves which underlie arrhythmia. Conversely, blocking hyper-phosphorylation of RyRs reverses channel dispersion in HF cells and associated impairment of Ca^2+^ release, providing new insight into the protective mechanisms of β-blockade in this disease.

## Results

### Prolonged β-adrenergic receptor (β-AR) activation causes RyR cluster dispersion

Using 3D dSTORM imaging, we investigated the effects of long-term phosphorylation on RyR organization in cardiomyocytes. To this end, we compared RyR arrangements in normal rat ventricular cardiomyocytes to those treated with isoproterenol (100 nM) for varying lengths of time (10, 30 or 60 min). 3D dSTORM reconstructions revealed progressive RyR cluster fragmentation as the duration of isoproterenol application increased (Fig. 1a). Indeed, following a brief, 10 min bout of isoproterenol exposure, we detected a slight, but non-significant change in measured cluster parameters. However, after 30 minutes of isoproterenol treatment, RyR cluster size significantly decreased (Fig. 1b), while cluster density increased (Fig. 1d). By 1 hour of β-AR activation, continued RyR rearrangement was sufficient to reduce CRU size, as fewer RyRs remained within clusters located ≦100 nm from their neighbors (Fig. 1c, see enlargements in Fig. 1a for representative images of CRUs). Since overall RyR density was unchanged by isoproterenol treatment (Fig. 1e), the detected differences in RyR organization did not result from channel degradation, but rather indicate progressive dispersion of RyR clusters into more numerous, smaller groupings.

**Figure 1:**
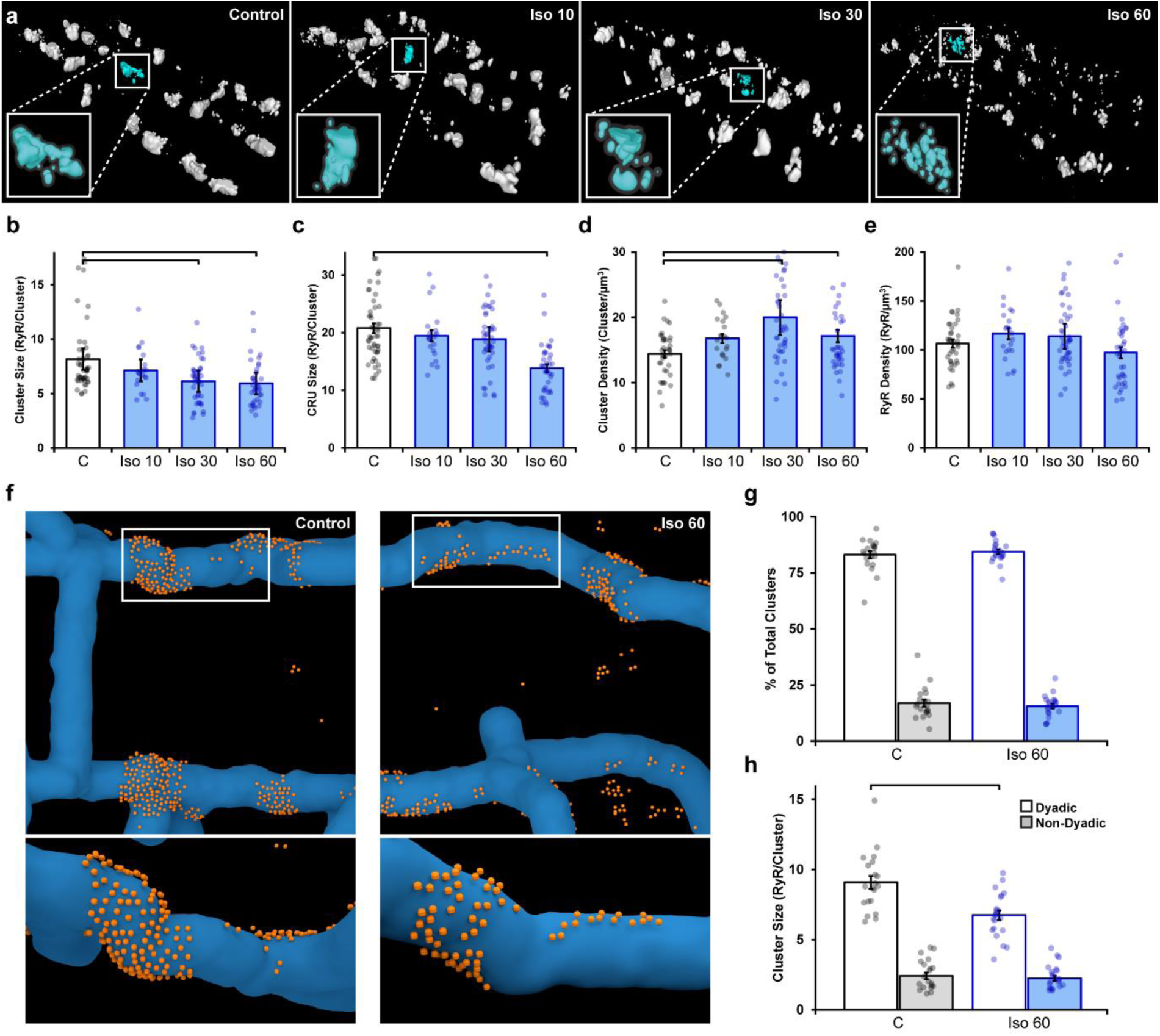
Prolonged β-adrenergic receptor (β-AR) activation disperses RyR clusters. (**a**) Representative reconstructions of internal RyR clusters based on 3D dSTORM in isolated rat cardiomyocytes. Control conditions are compared with isoproterenol treatment (100 nM) of varying duration (10, 30 and 60 minutes). The insets depict single CRUs, which encompass RyR clusters with edge-to-edge distances ≤ 100 nm. (**b-d**) Quantification of the dSTORM data revealed progressive dispersion of RyR clusters during isoproterenol treatment, as indicated by measurements of RyR cluster size, CRU size, and cluster density. (**e**) RyR density was unchanged. (**Control**: n_cells_ = 50, n_hearts_ = 6; **Iso 10**: n_cells_ = 21, n_hearts_ = 3; **Iso 30**: n_cells_ = 43, n_hearts_ = 4; **Iso 60**: n_cells_ = 37, n_hearts_ = 5). (**f**) Correlative imaging of the transverse tubular network (confocal microscopy) and RyRs (dSTORM) was employed to create 3D reconstructions of dyadic and non-dyadic CRUs (T-tubules: blue; RyRs: orange). (**g**) Similar proportions of clusters were characterized as dyadic under control and following 60 min isoproterenol. (**h**) Cluster size measurements indicated that dispersion during isoproterenol occurred exclusively within dyads (Control: n_cells_ = 20, n_hearts_ = 3; ISO 60: n_cells_ = 21, n_hearts_ = 3).

We next performed 3D correlative imaging of RyRs and the t-tubular network to distinguish between dyadic and non-dyadic (“orphaned”) RyRs (Fig. 1f). Based on these analyses, the majority of RyR clusters were classified as dyadic, and this proportion was observed to be similar under control and isoproterenol-treatment conditions (83 ± 1.6% and 84 ± 1.0%, respectively; Fig. 1g). Importantly, RyR cluster dispersion was found to be restricted to RyRs within dyads (control: 9.0 ± 0.5 vs 60 min isoproterenol: 6.7 ± 0.4 RyRs/cluster, Fig. 1h). It should be noted that the above analyses were performed on RyR clusters within the cell interior. Interestingly, the organization of surface RyR clusters associated with the cardiomyocyte sarcolemma was unaffected by β-AR activation (Supplementary Fig. 1), suggesting that RyR dispersion is a phenomenon restricted to internal, dyadic RyRs. We thus focused only on interior RyR clusters for the remainder of the study.

### Ca^2+^/calmodulin-dependent protein kinase II (CaMKII) activation mediates isoproterenol-induced RyR cluster dispersion

In cardiomyocytes, it is well established that a substantial portion of β-AR signaling is mediated via CaMKII^18^, and includes CaMKII-mediated phosphorylation of RyRs^19,20^. Indeed, using flow cytometry analysis, we observed increased RyR phosphorylation at the CaMKII site ser-2814, that was particularly prominent during early stages of isoproterenol treatment (Supplementary Fig. 2a-c). We therefore hypothesized that isoproterenol-induced cluster dispersion is, at least in part, driven by the actions of CaMKII. To this end, we first tested effects of the CaMKII competitive inhibitor autocamtide-2-related inhibitory peptide (AIP, 2 μM) following an initial hour-long isoproterenol incubation (i.e., 1 h isoproterenol followed by 1 h isoproterenol + AIP). We indeed found that the addition of AIP markedly reversed cluster dispersion (Fig. 2a). This was reflected by a significant increase in both RyR cluster size (Fig. 2d) and CRU size (Fig. 2e), as well as a significant reduction in RyR cluster density (Fig. 2f). Furthermore, direct activation of CaMKII through the Epac2 pathway using 8-CPT-cAMP (10 μM)^21^ consistently reproduced the RyR cluster dispersion observed during isoproterenol stimulation (Fig. 2b, mean measurements in Fig. 2d-f). RyR dispersion was again reversed by the addition of AIP post-treatment in these cells. To further corroborate our results, we indirectly stimulated CaMKII activity by applying a low concentration of caffeine (0.5 mM) to cardiomyocytes to increase spontaneous Ca^2+^ release from the SR^22^. We found that caffeine application gave rise to significant reductions in both RyR cluster and CRU size, as well as a significant increase in cluster density (Fig. 2c, mean data in Fig. 2d-f). Again, these changes were reversed when AIP was administered.

**Figure 2:**
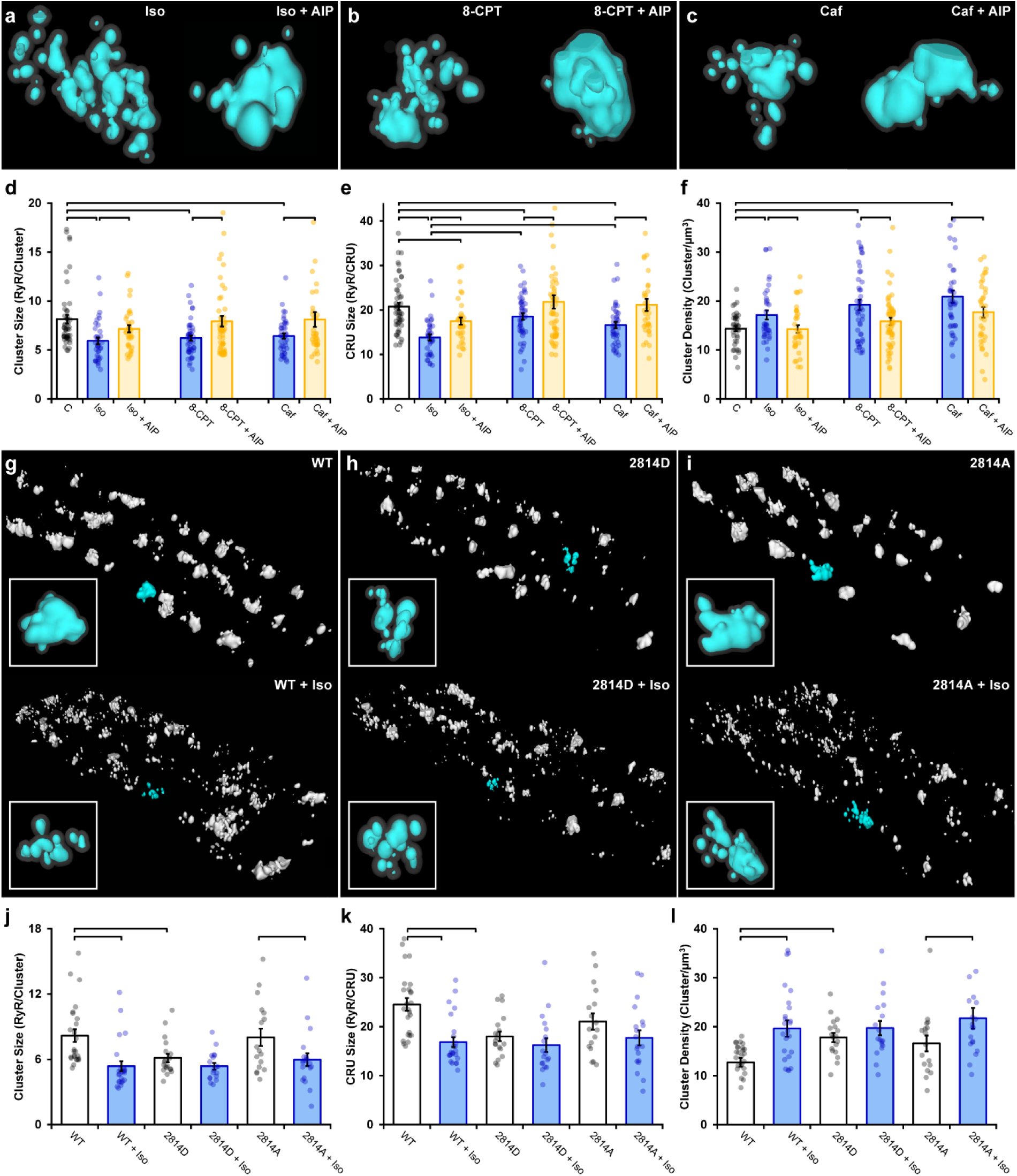
Ca^2+^/calmodulin-dependent protein kinase II (CaMKII) activation promotes RyR cluster dispersion. (**a-c**) Representative CRUs imaged in cardiomyocytes treated with isoproterenol (100 nM), 8-CPT (10 μM), or caffeine (0.5 mM) for 1 hour (left panels) or with with inclusion of the CaMKII inhibitor AIP (2 μM) for an additional hour (right panels). (**d-f**) Induced RyR cluster dispersion was reversed by CaMKII inhibition, as indicated by measurements of RyR cluster size, CRU size, and cluster density. (**Control**: n_cells_ = 50, n_hearts_ = 6; **Iso**: n_cells_ = 37, n_hearts_ = 5; **Iso + AIP**: n_cells_ = 37, n_hearts_ = 5; **8-CPT**: n_cells_ = 48, n_hearts_ = 5; **8-CPT + AIP**: n_cells_ = 52, n_hearts_ = 5; **Caffeine**: n_cells_ = 42, n_hearts_ =4; **Caffeine + AIP**: n_cells_ = 35, n_hearts_ = 3). **(g-h)** Representative images of RyR organization in cardiomyocytes from wild type mice (WT) and transgenic mice constitutively activated (S2814D) or genetically ablated (S2814A) phosphorylation at S2814. Images and mean data (**j-l**) are presented under baseline conditions and following 60 min isoproterenol stimulation. (**WT**: n_cells_ = 25, n_hearts_ = 2; **WT + Iso**: n_cells_ = 24, n_hearts_ = 2; **S2814D**: n_cells_ = 19, n_hearts_ = 2; **S2814D + Iso**: n_cells_ = 18, n_hearts_ = 2; **S2814A**: n_cells_ = 17, n_hearts_ = 2; **S2814A+ Iso**: n_cells_ = 19, n_hearts_ = 2).

While these findings suggest that isoproterenol-induced cluster dispersion is dependent on CaMKII, it remained unclear whether these effects can be directly attributed to its phosphorylation of RyRs. To address this issue, we examined RyR cluster characteristics in cardiomyocytes isolated from transgenic mice in which the CaMKII phosphorylation site on RyR is either constitutively active (S2814D) or genetically ablated (S2814A). We found that the RyR arrangement in the phosphomimetic S2814D mutant cells *at baseline* resembled the fragmented RyR organization observed in wild type (WT) animals after isoproterenol treatment (Fig. 2g, h). Indeed, cluster size (Fig. 2j), CRU size (Fig. 2k), and cluster density (Fig. 2l) were all similar in WT + ISO and S2814D cardiomyocytes, and significantly different from untreated WT cells. Furthermore, application of isoproterenol to S2814D cardiomyocytes did not lead to significant additional RyR dispersion, and only tendencies toward smaller cluster and CRU sizes (Fig. 2h, j-l). In contrast, cardiomyocytes from non-phosphorylatable S2814A mutant mice exhibited similar RyR organization as in WT mice at baseline (Fig. 2i). Administering isoproterenol to this group promoted a significant reduction in cluster size (Fig. 2j) and increased cluster density (Fig. 2l). These findings mirror observations in rat cardiomyocytes, where CaMKII inhibition did not fully reverse RyR dispersion in several experiments (Fig. 2d-f). Taken together, these results support that CaMKII-dependent RyR phosphorylation during prolonged β-AR stimulation is a key driver for cluster dispersion, but that there are likely additional contributing mechanisms.

### Protein Kinase A (PKA)-dependent phosphorylation also contributes to RyR cluster dispersion during β-AR activation

Following β-AR stimulation, increased cyclic AMP levels lead to PKA-dependent phosphorylation of the RyR at ser-2808^17^ (Supplementary Fig. 2d). Based on our above results, we hypothesized that in parallel to CaMKII-mediated RyR phosphorylation, activation of PKA following isoproterenol application also contributes to RyR cluster dispersion. Indeed, β-AR-induced cluster dispersion was reversed by the addition of the PKA inhibitor H89 (Fig. 3a), as reflected by significantly increased cluster size (Fig. 3c) and CRU size (Fig. 3d), and reduced cluster density (Fig. 3e). To further validate an involvement of PKA, we quantified RyR arrangement following application of the selective PKA activator 6MB-cAMP (6MB, 100 μM)^23,24^. As illustrated in Fig. 3b, 6MB-cAMP treatment prompted RyR dispersion comparable to that elicited by isoproterenol. These changes were again reversible by co-incubating 6MB with H89 following an initial treatment period with 6MB alone (Fig. 3c-e). As in experiments investigating effects of CaMKII inhibition (Fig. 2), it should be noted that PKA inhibition did not fully restore the RyR configuration at the CRU level. These data support that CaMKII- and PKA-dependent phosphorylation of RyRs concertedly drive RyR dispersion during prolonged β-AR stimulation.

**Figure 3:**
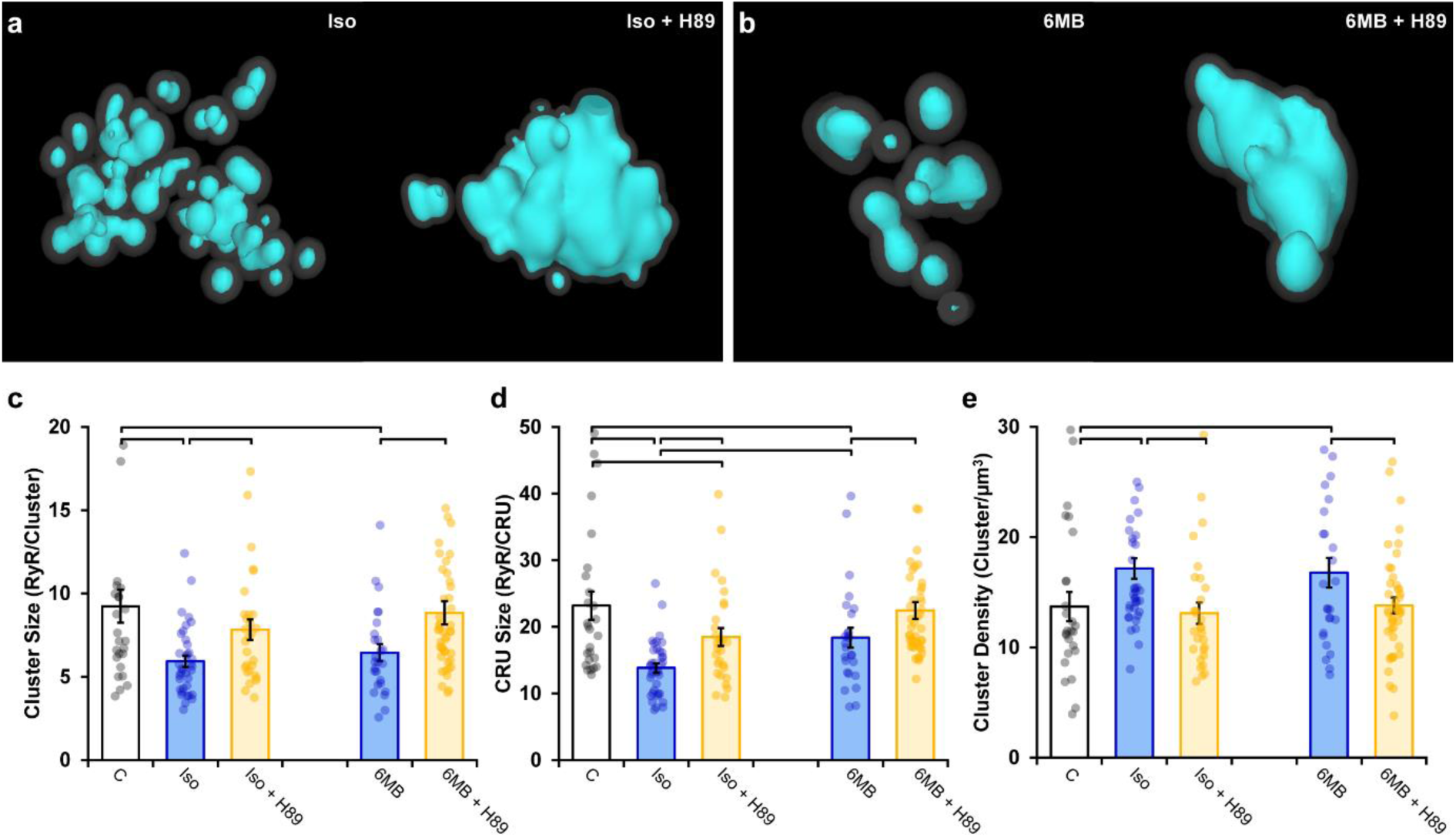
Protein kinase A (PKA)-dependent phosphorylation also promotes RyR cluster fragmentation during β-AR activation. **(a)** Representative CRUs indicate that PKA inhibition (H89, 10 μM) reverses isoproterenol-induced cluster dispersion. (**b**) Direct activation of PKA using 6MB-cAMP (100 μM) can also induce cluster dispersion, and is similarly reversed when followed by H89 application. (**c-e**) Mean data comparing cluster size, CRU size and cluster density between groups (Control: n_cells_ = 26, n_hearts_ = 3; Iso: n_cells_ = 37, n_hearts_ = 5; Iso + H89: n_cells_ = 30, n_hearts_ = 4; 6MB: n_cells_ = 26, n_hearts_ = 3; 6MB + H89: n_cells_ = 44, n_hearts_ = 4).

### β-stimulation-induced RyR dispersion alters Ca^2+^ sparks, transients, and waves

We next assessed the functional consequences of RyR dispersion. We examined spontaneous Ca^2+^ sparks since these events occur almost exclusively at dyads^25^, where RyR arrangements appear to be sensitive to β-AR stimulation (Fig. 1f). Previous work has shown that acute β-AR activation increases the frequency of spontaneous Ca^2+^ sparks, while diverse effects on spark geometry are reported^26-28^. We hypothesized, however, that RyR dispersion during prolonged β-AR stimulation would slow spark kinetics and increase Ca^2+^ leak, as released Ca^2+^ propagates between multiple clusters. Representative Ca^2+^ spark recordings from control and isoproterenol-treated cells are presented in Fig. 4a and b, respectively, with corresponding spark time courses presented in the right panels. We observed that while spark frequency was augmented throughout the isoproterenol treatment period (Fig. 4c), there was indeed a progressive slowing of both spark rise time (Fig. 4d) and duration (Fig. 4e). Increased overall spark geometry (Fig. 4f, g) during prolonged isoproterenol treatment was linked to progressively augmented Ca^2+^ leak, as calculated based on spark mass and frequency (Fig. 4h). Importantly, reversal of RyR dispersion by co-incubation of cells with either AIP or H89 after the initial isoproterenol treatment also reversed slowing of spark kinetics and isoproterenol-induced Ca^2+^ leak (Fig. 4d-h).

**Figure 4:**
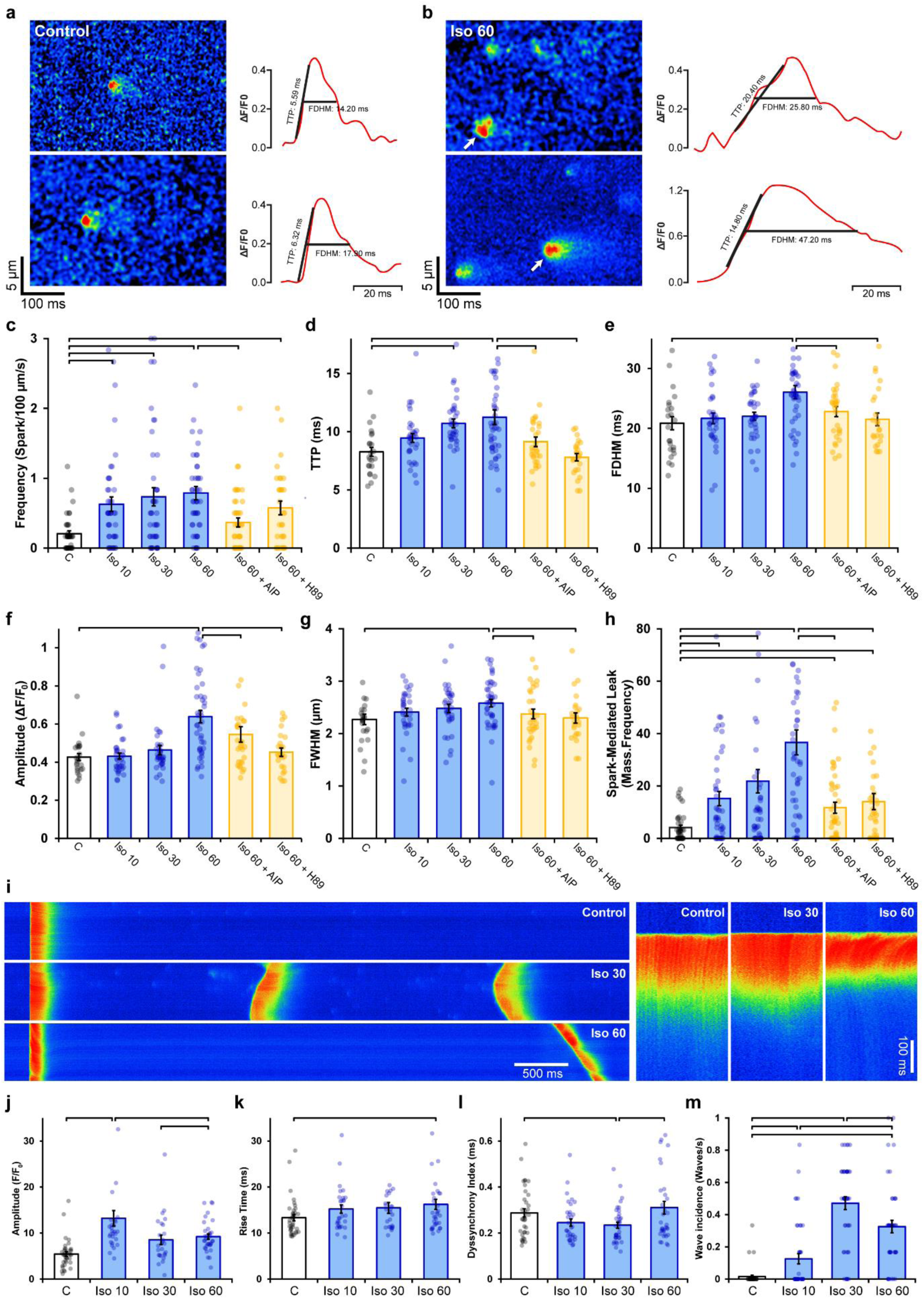
Effects of RyR dispersion of intracellular Ca^2+^ homeostasis are reversed by CaMKII or PKA inhibition. **(a, b)** Representative examples of Ca^2+^ sparks of control and isoproterenol (60 min)-treated cardiomyocytes obtained by confocal line-scan imaging. The time course for each spark is shown at right, with indicated TTP and FDHM measurements. (**c-h**) Mean data indicate that increasing spark frequency, geometry, and slowing of kinetics during isoproterenol were all reversed by CaMKII or PKA inhibition (Control: n_cells_ = 39, n_hearts_ = 3; Iso 10: n_cells_ = 44, n_hearts_ = 3; Iso 30: n_cells_ = 44, n_hearts_ = 3; Iso 60:n_cells_ = 47, n_hearts_ = 3; Iso 60 + AIP: n_cells_ = 45, n_hearts_ = 3; Iso 60 + H89 n_cells_ = 31, n_hearts_ = 2). (**i**) Representative linescan images illustrating the last in a series of electrically-paced Ca^2+^ transients, followed by a pause to examine Ca^2+^ wave generation. Enlargements of the Ca^2+^ transients are shown at right to highlight differences in Ca^2+^ release synchrony. (**j-l**) Mean data showed that initial increases in Ca^2+^ transient amplitude and synchrony were reversed with continued exposure to isoproterenol, while overall release kinetics slowed. (Control: n_cells_ = 38, n_hearts_ = 3; Iso 10: n_cells_ = 35, n_hearts_ = 3; Iso 30: n_cells_ = 36, n_hearts_ = 3; Iso 60: n_cells_ = 41, n_hearts_ = 3). (**m**) Ca^2+^ wave incidence increased during early time points following isoproterenol treatment, but then reversed. (Control: n_cells_ = 43, n_hearts_ = 3; Iso 10: n_cells_ = 44, n_hearts_ = 3; Iso 30: n_cells_ = 46, n_hearts_ = 3; Iso 60:n_cells_ = 44, n_hearts_ = 3).

Changes in RyR localization and function during prolonged β-AR stimulation were additionally linked to alterations in cell-wide Ca^2+^ transients (Fig. 4i). As expected, isoproterenol treatment acutely increased Ca^2+^ transient magnitude, however this reversed with continued exposure (Fig. 4j). This pattern was mirrored by changes in SR Ca^2+^ content (Supplementary Fig. 3), which was also initially increased by heightened SERCA activity during isoproterenol exposure, but then began to decline as RyR Ca^2+^ leak progressively increased. Similar changes were observed in the synchrony of Ca^2+^ release across the cell. Ca^2+^ release tended to be more synchronous at early stages of isoproterenol treatment (Fig. 4l), in agreement with previous work indicating that increasing SR Ca^2+^ content and RyR sensitization enable more homogeneous Ca^2+^ transients^22,29^. However, after prolonged β-AR stimulation, RyR dispersal, slowed Ca^2+^ spark kinetics, and declining SR content were coupled to a reversal of the synchronization of Ca^2+^ release observed at earlier time points (Fig. 4l), and a modest slowing of the rising phase of the transient (Fig. 4k).

The above data are consistent with the notion that RyR dispersion gradually counters the stimulatory effects of β-AR stimulation on CICR, perhaps in a protective manner. To further examine this hypothesis, we investigated the propensity for spontaneous Ca^2+^ waves during isoproterenol treatment (Fig. 4i). Indeed, while we observed an expected initial increase in Ca^2+^ wave incidence following isoproterenol application, this reversed after 60 min as wave frequency was reduced by ∼30% (Fig. 4m). Thus, sufficient dispersion of RyRs and accompanying weakening of CICR inhibits wave propagation, suggesting a protective role or RyR rearrangement during prolonged β-AR stimulation.

### RyR cluster dispersion lowers CRU excitability in a mathematical model

To support the above correlative data linking RyR arrangement and Ca^2+^ homeostasis, we employed a mathematical reaction-diffusion model of Ca^2+^ release in the CRU. Using paired images of RyRs (3D dSTORM) and t-tubules (confocal microscopy), we generated four computational CRU geometries that were representative of the range of RyR dispersion observed in cells (Fig. 5a, Supplementary Table 1). For each geometry, a series of 400 stochastic spark simulations was performed with and without the regulatory effects of β-AR stimulation on RyR sensitivity and SR load included (Fig. 5b). Other model parameters are described in Supplementary Tables 2 and 3.

**Figure 5.**
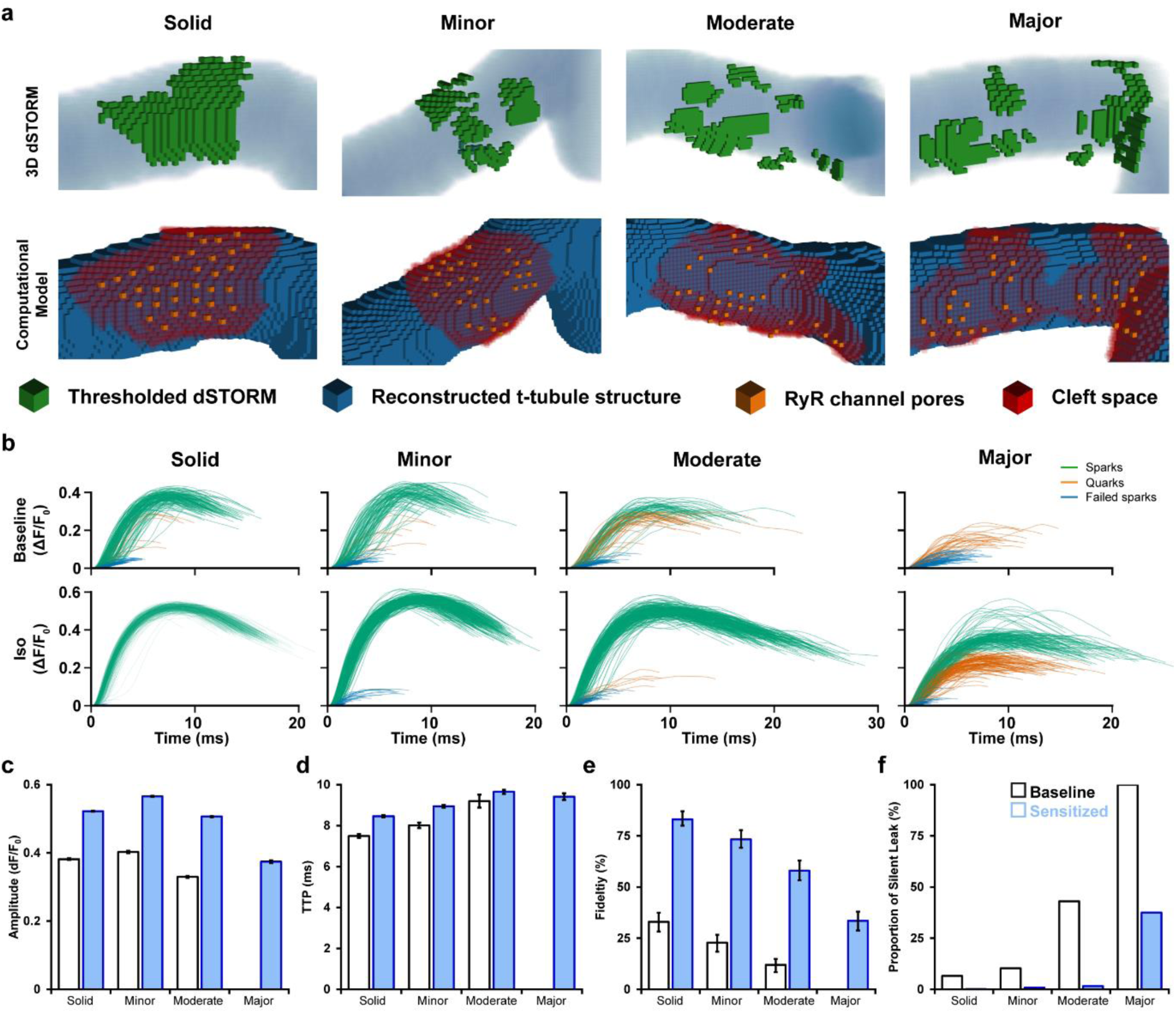
Mathematical modeling linking changes in RyR configuration and function during β-adrenergic stimulation. (**a**) 3D computational volumetric meshes built from correlative imaging of the t-tubular network (confocal microscopy) and RyRs (3D dSTORM). Constructed geometries consisted of 10×10×10 nm voxels and a 1×1×1 μm volume. The four geometries were chosen to represent the range of cluster dispersion seen in imaging, see Table 1 for details. (**Top row**) Reconstructed t-tubular structures (blue) are illustrated with superimposed thresholded RyR signals (green). (**Bottom Row**) Corresponding computational geometries, with indicated RyR channel pores (orange) and cleft space (red) defined by morphological dilation around the RyRs. The four geometries were selected to represent the range of RyR dispersion observed in experiments, ranging from solid to majorly-fragmented CRUs (see methods). (**b**) 400 stochastic spark simulations were performed in each geometry with baseline model parameters, and another 400 were conducted with the regulatory effects of β-AR stimulation added (RyR sensitization and increased SR content). The ΔF/F_0_ time course is shown for each spark. The maximal amplitude used to define the release as an experimentally observable spark-event (ΔF/F_0_ ≥ 0.3), a sub-spark quark event (ΔF/F_0_ ≥ 0.1), or a failed spark (ΔF/F_0_ < 0.1). **(c, d)** Mean amplitude and time to peak (TTP) measurements for observable sparks. (**e, f**) The classification of release into spark and non-spark events was used to estimate spark fidelity and the ratio of leak that is “silent” (i.e., sub-spark release) in each simulation case. Error bars in C and D show standard error, while the bars in E indicate the 95% Agresti-Coull confidence interval.

Each simulated Ca^2+^ release event was classified as being an observable spark event (ΔF/F_0_ ≥ 0.3), a sub-spark event (Ca^2+^ quark^30^), or a failed spark for which negligible Ca^2+^ release occurred (ΔF/F_0_ ≤ 0.1; Fig. 5b). The proportion of observable sparks elicited was used as an estimate of spark fidelity (Fig. 5e). With baseline model parameters, RyR cluster dispersion was observed to have a large impact on CRU excitability. Indeed, spark fidelity declined steadily with increased fragmentation (95% CI in solid CRU: 28.6-37.8% vs majorly fragmented CRU: 0-1.2%, Fig. 5e). However, β-AR stimulation augments both RyR sensitivity and, particularly at early stages, SR Ca^2+^ content. Increasing channel sensitivity and Ca^2+^ release flux accordingly in the model improved inter-RyR communication, and increased spark fidelity. In the cases of the minorly- and moderately-dispersed CRUs, these effects more than compensated for the effects of RyR fragmentation, as spark fidelity even exceeded values obtained in solid CRUs at baseline (Fig. 5b, e). However, CRUs with greater degrees of RyR dispersion continued to exhibit lower fidelity of spark generation than those with more compact arrangements.

Analyzing only successful spark events revealed that RyR cluster dispersion also influenced spark amplitude (Fig. 5c) and kinetics (Fig. 5d). Individual simulation results are presented alongside experimental data in Supplementary Fig. 4. While minor to moderate degrees of RyR dispersion had rather little impact on spark amplitude, majorly fragmented CRUs exhibited markedly smaller events (solid w/β-AR: ΔF/F_0_ = 0.52 ± 0.001, majorly fragmented w/β-AR: ΔF/F_0_ = 0.37 ± 0.004, Fig. 5c). This decrease was attributed to a shift from an all-or-none pattern of RyR activation in solid CRU arrangements to partial activation in fragmented units (Fig. 5b). The time to peak (TTP) of sparks increased with the degree of dispersion in our model (Fig. 5d) and, under simulated β-AR activation, this was primarily due to the sensitized RyRs’ improved ability to sustain regenerative release between clusters. Indeed, in simulations where SR load was systematically altered with constant RyR Ca^2+^ sensitivity, spark kinetics were not markedly affected (Supplementary Fig. 5). These findings support that RyR cluster dispersion during prolonged β-AR activation is directly linked with slower, and lower magnitude Ca^2+^ sparks observed experimentally.

From the classification of each simulation as either a spark or sub-spark event, we also calculated the proportion of total RyR-mediated Ca^2+^ leak that is “silent”, rather than spark-mediated (Fig. 5f). Due to declining spark fidelity, we observed that although total Ca^2+^ released from a given CRU decreased with greater fragmentation, the proportion of leak which is silent increased dramatically. When including the effects of β-AR stimulation, fidelity increased and the ratio of silent leak was strongly reduced, becoming close to zero for the mildly to moderately fragmented CRUs. For the majorly fragmented CRU, however, there remained a considerable amount of silent leak (Majorly fragmented w/β-AR: 37.5% silent leak, Fig. 5f) linked to partial activation of the CRU (Ca^2+^ quarks).

Taken together, the modeling results indicate that RyR Ca^2+^ leak shifts from spark-mediated to silent leak when CRUs are dispersed, reducing the fidelity of spark generation. Thus, while RyR phosphorylation during acute β-AR stimulation markedly enhances spark fidelity, gradual CRU fragmentation during continued stimulation reverses these changes. An accompanying reduction in SR content is expected to exacerbate declining spark fidelity during prolonged β-AR stimulation. Furthermore, when sparks are successfully elicited within dispersed CRU geometries, Ca^2+^ release is slowed and of lower magnitude.

### CaMKII and PKA activation promote RyR dispersion and Ca^2+^ dysregulation during heart failure (HF)

We next examined whether dispersion of RyRs reported during HF^11^ similarly results from RyR hyper-phosphorylation. Previous work has indeed linked CaMKII^15,16^ and PKA activity^17^ to RyR dysfunction in this disease. We first examined RyR phosphorylation status in rats with post-infarction HF compared to sham-operated controls (see Supplementary Table 4 for animal characteristics). Western blotting revealed no significant change in total RyR expression, and only a tendency toward increased PKA phosphorylation at S2808 (Fig. 6a, b – source data 1). However, phosphorylation at S2814 - a site linked to CaMKII activation - was significantly increased by ∼55% in HF (Fig. 6b – source data 1).

**Figure 6.**
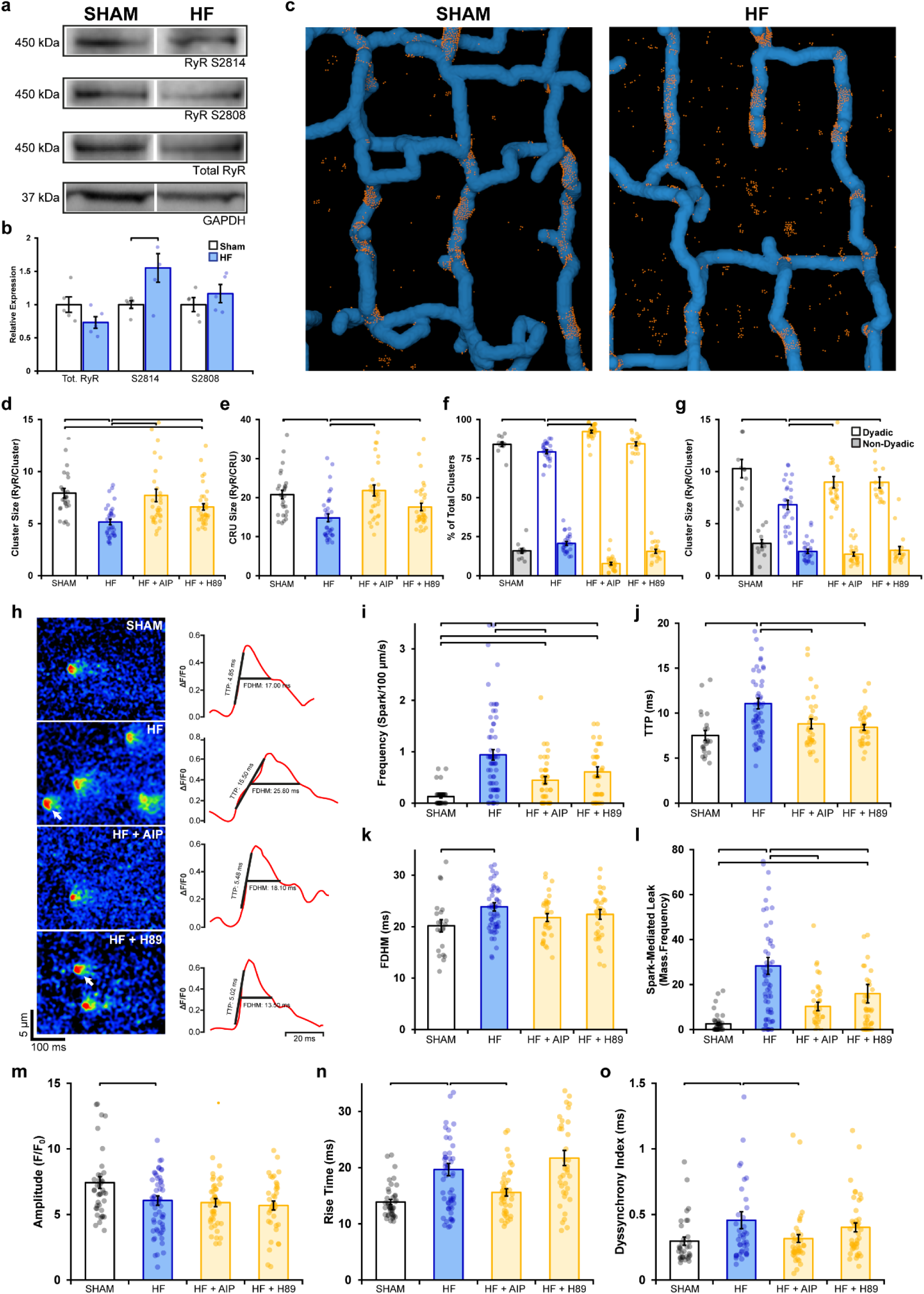
Remodeling of RyR organization and function during heart failure (HF) are reversed by inhibiting phosphorylation. **(a, b)** Representative Western blots and mean data comparing the relative expression of CaMKII-phosphorylated RyR (S2814), PKA-phosphorylated RyR (S2808) and total RyR in Sham and HF cardiomyocytes. (Sham: n_hearts_ = 5; HF: n_hearts_ = 5). (**c**) 3D reconstruction of dyads in Sham and failing cells based on correlative imaging of t-tubules (blue, confocal microscopy) and RyRs (orange, 3D dSTORM). (**d-g**) Mean RyR cluster size, CRU size, and RyR cluster density in cardiomyocytes from Sham and HF cells, at baseline and following inhibition of CaMKII (AIP) or PKA (H89). (Sham: n_cells_ = 27, n_hearts_ = 4; HF: n_cells_ = 33, n_hearts_ = 4; HF + AIP: n_cells_ = 33, n_hearts_ = 4; HF + H89: n_cells_ = 35, n_hearts_ = 4). (**h**) Representative Ca^2+^ spark examples from Sham and HF cardiomyocytes with or without CaMKII inhibition (AIP) or PKA inhibition (H89). The time course of each spark is shown at right. (**i-l**) Mean data comparing spark duration, time to peak, frequency and spark-mediated leak in Sham and HF cells, at baseline and following treatment with AIP or PKA (Sham: n_cells_ = 42, n_hearts_ = 3; HF: n_cells_ = 65, n_hearts_ = 5; HF + AIP: n_cells_ = 63, n_hearts_ = 5; HF + H89: n_cells_ = 43, n_hearts_ = 3). (**m-o**) Mean measurements of cell-wide Ca^2+^ transient magnitude, TTP (rise time), and dyssynchrony in Sham and HF cells, in the presence and absence of AIP or PKA (Sham: n_cells_ = 41, n_hearts_ = 4; HF: n_cells_ = 59, n_hearts_ = 4; HF + AIP: n_cells_ = 43, n_hearts_ = 4; HF + H89: n_cells_ = 38, n_hearts_ = 3).

3D dSTORM imaging of Sham and HF cardiomyocytes (Fig. 6c) revealed significant RyR dispersion in failing cells, manifested at both the cluster level (Fig. 6d; Sham: 7.93 ± 0.46 vs HF: 5.16 ± 0.27 RyRs/Cluster) and CRU level (Fig. 6e; Sham: 20.79 ± 1.09 vs HF: 14.82 ± 1.01 RyRs/CRU). These structural changes are consistent with a previous 2D super-resolution study which utilised the same pathological model^11^. Next, we subjected cells to either CaMKII or PKA inhibition to assess whether disrupted RyR organisation could be reversed. Indeed, inhibiting CaMKII phosphorylation with AIP in HF cells restored RyR organization to Sham values, as evidenced by a 49% increase in cluster size and a 47% increase in CRU size (Fig. 6d, e). To a lesser extent, the addition of H89 to failing cells also significantly reversed RyR dispersion at the cluster level (28% increase) and CRU level (19% increase). Correlative imaging of RyRs and T-tubules (Fig. 6c) revealed a significantly smaller proportion of dyadic clusters in HF compared to Sham (Fig. 6f); an expected finding since reduced t-tubule density in this model results in the formation of “orphaned” RyRs^31,32^. The smaller RyR clusters observed in failing cells were found to exclusively occur at dyadic sites (Sham: 9.09 ± 0.47 vs HF: 6.81 ± 0.44 RyRs/cluster), where dispersed RyR arrangements were again reversible by CaMKII or PKA inhibition (Fig. 6g). The striking similarity of these findings to experiments employing prolonged isoproterenol exposure (Figs. 1-4) supports that fragmentation of RyR clusters during HF results from prolonged β-AR activation stimulation in this condition, linked to the activation of both CaMKII and PKA.

Finally, we examined whether alterations in RyR Ca^2+^ release during HF could be linked to RyR phosphorylation and dispersion. Representative recordings of Ca^2+^ sparks are presented in Fig. 6h, together with temporal profiles shown at right. Failing cardiomyocytes exhibited significantly increased frequency (Fig. 6i), slower rise time (Fig. 6j), and longer duration sparks (Fig. 6k) than in sham cells, which summated to marked augmentation of spark-mediated leak (Fig. 6l). All of these changes were reversed by inhibiting CaMKII or PKA, although the effects of CaMKII inhibition were the most marked (Fig. 6h-l). Overall Ca^2+^ transients were of lower magnitude in failing cardiomyocytes (Fig. 6m), and Ca^2+^ release was observed to be slowed and dyssynchronous (Fig. 6n, 6o, see Supplementary Fig. 6 for representative examples). CaMKII inhibition again reversed changes in Ca^2+^ transient kinetics and synchrony in failing cells to values similar to Sham (Fig. 6n, o). Taken together, these findings identify phosphorylation-dependent RyR dispersion as a key driver of RyR dysfunction in HF, and particularly implicate CaMKII overactivity in this process.

## Discussion

In this work, we identify an important new mechanism by which RyR organization and function are controlled by β-AR stimulation. Specifically, we show that during prolonged β-AR activation, as occurs during HF, there is a progressive dispersion of RyR clusters. This fragmentation is driven by both CaMKII- and PKA-dependent phosphorylation of the channel, and is reversible by inhibiting these kinases. Functionally, our experimental and modeling results indicate that dispersed CRUs exhibit increased “silent” Ca^2+^ leak, and low spark fidelity. When Ca^2+^ sparks are successfully generated by dispersed CRUs, they exhibit slowed kinetics and reduced magnitude. These changes are in turn coupled with smaller and desynchronized Ca^2+^ transients, which are hallmarks of failing cells. However, we also observed protective actions of CRU dispersion, as lowered spark fidelity and magnitude protected against Ca^2+^ wave generation. Thus, RyR dispersal appears to be aimed at curbing arrhythmogenesis during prolonged β-AR stimulation, but comes at the expense of systolic function, and this is particularly detrimental in the setting of HF.

While the precise nature of inter-RyR dynamics remains incompletely understood, several previous studies have suggested that there are important structural determinants of CRU function^11,33-35^. Of note, recent work by Galice *et al*. (2018) measured RyR cluster size and Ca^2+^ sparks simultaneously, finding that spark frequency depends steeply on cluster size, while spark amplitudes appear to be independent of size above a certain threshold. The authors argued that RyR cluster sizes must strike a balance between being sufficiently large to ensure high-fidelity CICR, but not so large as to promote arrhythmogenic behavior. When RyR channels are sensitized or Ca^2+^ levels are elevated, this balance shifts, increasing the propensity for sparks, diastolic leak and waves. Extending from these findings, our present experimental and mathematical modeling results suggest that RyR cluster dispersion compensates for this shift by physically inhibiting Ca^2+^ wave propagation, but to the detriment of spark fidelity.

Our data show that marked RyR cluster dispersion occurs only after prolonged β-AR stimulation, which likely explains why previous, acute-treatment studies have not detected similar changes in RyR arrangement or function. For example, Asghari *et al*.^14^ recently performed detailed analysis of RyR positioning, but within only 10 minutes of channel phosphorylation. Nevertheless, these authors did observe changes in the precise positioning of neighboring RyRs within this timeframe, as a checkerboard arrangement of RyRs was favored over side-by-side arrangements. Both arrangements involve SRPY and P1 domain-domain interactions between adjacent receptors^36^. A shift in the configuration of these domains, for example during phosphorylation, might be envisioned to reduce the stability of RyR dimerization, i.e., reducing RyR “stickiness”. Indeed, Asghari *et al*^14^ observed a modest rightward shift in the RyR-RyR nearest neighbor distance early after initiation of phosphorylation, which could represent the earliest stages of cluster breakup that we have observed at later time points. However, neither we (Supplementary Fig. 1) nor Asghari *et al* observed similar rearrangement of RyRs at the cell surface during β-AR stimulation, suggesting that there are unique features of internal RyRs and/or their partner proteins that render their positioning phosphorylation-sensitive. FKBP12.6 is one such candidate, as it has been shown to be directly involved in domain-domain interactions between adjacent RyRs^36^, and to fine-tune RyR positions in directions which are opposite those of phosphorylation^14^.

It is well established that prolonged β-AR stimulation is linked to time-dependent desensitization of the signaling pathway; a process that involves G protein-coupled receptor kinases and β-arrestins which promote internalization and downregulation of the receptors^37^. Our results indicate that this process was likely initiated in our experiments, since RyR phosphorylation at ser-2814 and ser-2808 both exhibited bell-shaped responses during the 1 h treatment protocol (Supplementary Fig. 2). A similar response was observed in measurements of SERCA activity (Supplementary Fig. 3c). Importantly, RyR phosphorylation by CaMKII and/or PKA remained elevated above control levels at the end of the protocol. These heightened phosphorylation levels apparently remain sufficient to support RyR dispersion, since inhibition of either kinase following β-AR stimulation caused RyRs to re-cluster (Fig. 2a-f, 3). The regulatory role of CaMKII and PKA on RyR dispersion was further corroborated by experiments showing that selective activation of either kinase induced fragmented CRU arrangements. Furthermore, transgenic mice with constitutively active S2814 were found to have significantly dispersed clusters at baseline compared to their wild-type counterparts (Fig. 2g-l). Notably, genetic ablation of S2814 did not appear to inhibit the effects of β-AR stimulation, which continued to reduce RyR cluster size and increase cluster density. Taken together, these findings point to a more complicated mechanism responsible for cluster dispersion than direct phosphorylation of the RyR at the S2814 and S2808 residues by CaMKII and PKA, respectively. Rather, it seems likely that CaMKII and PKA act interdependently on the RyR. Other studies have also suggested cross-talk between the two sites, with one observing changes to S2814 phosphorylation in a S2808 mutant^38^, and another demonstrating increased PKA activity at S2808 following S2814 phosphorylation^39^. In HF cells, we detected an increase in RyR phosphorylation at S2814, but only a minor trend towards higher phosphorylation at S2808. Still, inhibition of either CaMKII or PKA reversed cluster dispersion in these cells, although the extent of reversion following AIP administration was more complete than that obtained with H89. These results appear to be consistent with studies suggesting that PKA may not be the principal determinant of RyR hyper-phosphorylation in HF^16,40^, but also indicate that CaMKII and PKA control RyR clustering in a coordinated manner.

Although we observed a close relationship between RyR organization and function in our experiments, we relied on mathematical modeling to directly interrogate the functional implications of cluster dispersion vs other regulatory effects of β-AR stimulation. The modeling results indicated that RyR cluster fragmentation is sufficient to significantly weaken intra-CRU RyR communication. Notably, these structural-driven changes in function were opposite those induced by regulatory changes, as increased RyR sensitivity to cytosolic Ca^2+^ and increased SR Ca^2+^ content increased intra-CRU RyR communication. These observations are consistent with the view that while acute β-AR stimulation effectively augments CICR, progressive RyR dispersion and accompanying decline of SR Ca^2+^ content during continued exposure serves to gradually reverse these changes, as Ca^2+^ release fidelity, magnitude, and kinetics are compromised (Fig. 5c-e). In healthy cardiomyocytes, this functional decline was not overly severe, as Ca^2+^ transients measured experimentally remained somewhat larger than in untreated cells, and Ca^2+^ release kinetics were only modestly slowed (Fig. 4j, k). However, during HF, more detrimental consequences of RyR dispersion are expected, as SR content is reduced below normal levels. Thus, compensatory changes in RyR function observed in healthy cells during β-AR stimulation are expected to be less present, and impairment of CICR more robust.

Experimental studies have frequently linked increases in spark frequency to higher incidence of Ca^2+^ waves^41,42^. Here, we also report a marked increase in both spark and wave incidence during the early stages of β-AR stimulation (Fig. 4i, m). However, while both spark frequency and mass continued to increase with prolonged treatment, wave incidence significantly declined. While this finding may at first seem counterintuitive, a similar ‘paradoxical’ observation was reported in a canine model of left ventricular hypertrophy^43^. The authors hypothesized that disconnects between junctional SR clusters and LTCCs could lead to spatial heterogeneity in SR Ca^2+^ content, enabling regions with lower SR Ca^2+^ to act as “fire breaks” against wave propagation. We suggest that an analogous mechanism explains our findings, but which may include both reduced SR content and dispersed CRUs acting as a physical barrier to Ca^2+^ wave propagation. Since accumulating evidence supports that RyR positioning is tightly regulated, we believe that RyR dispersion in healthy cardiomyocytes is, by design, aimed at curbing wave generation during prolonged β-AR stimulation. This is an important consideration if we are to consider targeting RyR localization in disease. While increasing RyR-RyR “stickiness” in failing cells would be expected to augment RyR clusters sizes, inter-channel collaboration, and systolic function, our data suggest that such effects may also increase susceptibility to arrhythmia.

In summary, we observed that RyR clusters progressively disperse during protracted β-AR activation, due to channel phosphorylation by both CaMKII and PKA. Experimental and mathematical modeling results linked this dyadic rearrangement to declining efficacy of CICR, including slowed and reduced magnitude Ca^2+^ release, but also protection against pro-arrhythmic Ca^2+^ waves. These findings have important implications for HF, and other conditions such as catecholaminergic polymorphic ventricular tachycardia, which are associated with increased phosphorylation of RyRs and risk of arrhythmia.

## Material and Methods

### Ethical approval

All animal experiments were performed in accordance with the Norwegian Animal Welfare Act and NIH Guidelines, and were approved by the Ethics Committee of the University of Oslo and the Norwegian animal welfare committee (FOTS ID 20208). The majority of the experiments were performed on adult male Wistar rats (250-350 g) purchased from Janvier Labs (Le Genest-Saint-Isle, France). Rats were group housed at 22°C on a 12 h:12 h light–dark cycle, with free access to food and water. Cardiomyocytes isolated from transgenic RyR2-S2814D and RyR2-2814A mice^44^ were provided by the laboratory of Xander Wehren (Baylor College of Medicine, Texas, United States). A total of 64 rats and 6 mice were used in this study. The authors understand the ethical principles under which *Nature Medicine* operates and declare that our work complies with this animal ethics checklist.

### Rat model of post-myocardial infarction congestive HF

Left coronary artery ligation was performed to induce large anterolateral myocardial infarctions in male Wistar rats^45^. Development of HF was verified six weeks later using a Vevo 2100 echocardiography imaging system (VisualSonics, Toronto, Canada). HF animals were selected based on established criteria^46^, including dilation of the left atrium (diameter >5 mm) and infarction size above 40% (Supplementary Table 4). Sham-operated rats served as controls. Sample sizes were determined by power analysis, assuming that only 50% of post-infarction animals would be included in the final data set, and based on a pilot project of variability in CRU morphology in healthy controls.

### Rat Ventricular Cardiomyocyte Isolation

Isolation of cardiomyocytes was based on the protocol previously described by Hodne *et al*.^*47*^. Briefly, animals were anaesthetized with isoflurane and sacrificed, and hearts were rapidly excised. Each heart was then cannulated and mounted on a constant-flow (3 ml/min) Langendorff setup, perfused with Ca^2+^-free oxygenated solution (in mmol/L: 140 NaCl, 5.4 KCl, 0.5 MgCl_2_, 0.4 NaH_2_PO_4_, 5 HEPES, 5.5 glucose pH 7.4). Once the heart had been cleared of blood, perfusion was switched to the same solution containing collagenase type II (1.8 mg/ml, Worthington Biochemical Corporation) for 10-12 mins at 37°C. Following digestion, left ventricular tissue was dissected and finely cut into 3-4 mm^3^ pieces. A secondary digestion was performed to liberate additional cells from tissue by transferring approximately 8 ml of tissue and collagenase solution to a 10 ml Falcon tube containing 0.2 mg DNase (LS002006, Worthington) in 500 μl BSA. Cells were subsequently filtered and allowed to pellet in 0.2 mmol/L Ca^2+^.

### Immunofluorescence Labeling

Isolated cardiomyocytes were washed with Dulbecco’s PBS (No. 4387, Biowhittaker), fixed with 4% PFA for 10 min, quenched with 100 μmol/L glycine for 10 min, and permeabilised with 1% triton X100 for 10 min. The cells were then plated on glass bottom dishes (No 1.5, Ø 14 mm, γ-irradiated, Martek Corporation) that had been coated with laminin (mouse, BD Biosciences) and blocked using Image-iT™ FX Signal Enhancer (Thermo Fisher Scientific) prior to immunolabeling.

For RyR visualisation, cells were incubated with mouse anti-RyR (1:100, MA3-916, Thermo Fisher Scientific) overnight at 4 °C. For t-tubule imaging, cells were incubated overnight at 4 °C in a mixture of rabbit anti-Cav-3 (1:100, ab2912, Abcam) antibody and a custom rabbit anti-NCX1 antibody as previously described^5^ (1:100, Genscript Corporation, Piscataway, NJ). Secondary antibody labeling was carried out using donkey anti-mouse Alexa Fluor 647 (1:200, A-21237, Thermo Fisher Scientific) and goat anti-rabbit Alexa Fluor 488 (1:200, A-11070, Thermo Fisher Scientific) antibodies for 1 h at room temperature. Both primary and secondary antibodies were diluted in a blocking buffer consisting of 2% goat serum and 0.02% NaN_3_ in PBS.

### 3D dSTORM Super Resolution Imaging and Reconstruction of RyRs

Alexa Fluor 647-labeled rat ventricular cardiomyocytes were submersed in an imaging buffer containing 20% VectaShield (H-1000, Vector Laboratories) diluted in TRIS-Glycerol (5% v/v TRIS 1 M pH 8 in Glycerol, Sigma Aldrich). This composition has been previously shown to produce comparable, if not superior quality dSTORM images when compared with conventional oxygen scavenging-dependent systems^48^.

Cardiomyocytes were imaged using the Zeiss ELYRA/LSM 710 system (Carl Zeiss; Jena, Germany). A diode laser (150 mW, 642 nm) illuminated the sample via a plan-apo 63x 1.4 NA oil objective, configured in a highly inclined and laminated optical sheet (HiLo). 3D imaging was achieved utilising Phase Ramp Imaging Localization Microscopy (PRILM) technology^49^. Fluorescence emission >655 nm was collected with an iXon 897 back-thinned EMCCD camera (Andor Technology, Belfast). A sequence of 15,000 frames was acquired for each cell at a frame exposure time of 40 ms. Throughout image acquisition, a piezo-operated Definite Focus system was employed to autocorrect for axial drift.

Reconstruction of dSTORM data was carried out using the ‘PALM Processing’ module in the ZEN Black software (Zeiss). In short, an experimental 3D PSF with an axial range of 4 μm was acquired using 100 nm TetraSpecks (T7279, Thermo Fisher Scientific). Individual single molecule events were detected by employing an 11 pixel circular mask, with a signal to background noise ratio of 6. Drift correction was performed using a 5 segment piece-wise linear function, and a text-based points table was then generated containing the x, y and z coordinates of each localization event. In order to minimize the inclusion of clusters with larger localization error, events from only the central 600 nm of the 4 μm stack were included^49^. Lastly, the points table was processed via a custom-written Python script, to generate a pixel based image whereby individual events were represented with a Gaussian function centered at the event coordinates and a width corresponding to its lateral and axial precision values (in nm). The resulting intensity data stack was then thresholded using the Ostsu method and output as a 600 nm z-stack with a voxel size of 30 nm, so that each voxel of the resulting 3D binary mask stack contained no more than a single RyR.

### Quantitative Analysis of 3D RyR Cluster Characteristics

3D quantification of RyR organization was performed as previously described^5^. In short, by combining Phase Ramp Imaging Localization Microscopy (PRILM) dSTORM and fluorescent event registration, we were able to discern the irregular shapes of internal RyR arrangements, and estimate RyR cluster and CRU sizes. Based on mathematical modeling by Sobie *et al*. ^3^, RyR clusters with edges localized within 100 nm were assigned to the same CRU. RyR cluster density was calculated and normalized by totaling the number of clusters within a 2 × 2 × 0.6 μm volume positioned within the cell interior. The volumetric density of RyRs was determined on a per cell basis by multiplying the average cluster size by the cluster density.

### Correlative Imaging of RyRs and t-tubules for Dyad Reconstruction

The Zeiss ELYRA system was also used for confocal imaging of RyRs (Alexa Fluor 647, 633 nm laser) and t-tubules (Alexa Fluor 488, 488 nm laser) for 3D correlative reconstruction of dyads. Prior to dSTORM imaging of RyRs, a 3 μM thick confocal stack centered on the same region was acquired (frame size: 1024 × 1024 pixels; pixel size: 50 nm; z spacing: 200 nm per slice). Deconvolution of confocal images was performed using Huygens Essential software (SVI, The Netherlands), with a signal to noise ratio of 5. Using the confocally-imaged RyR channel as a reference, confocally measured t-tubule distributions were correlated to dSTORM-derived RyR positions. This was done with a custom-written Python script which corrected for both lateral, axial, translational and scaling differences between the two imaging modalities to optimally align the dSTORM-derived RyR data with the confocal RyR data.

### 3D Geometric Rendering of RyRs and T-tubules

To examine the arrangement of RyR clusters in relation to t-tubules, 3D volumetric geometries were constructed from the correlated images of these structures using the same approach as described in our earlier work^5^. Geometries were built using a custom Python script relying on NumPy 1.18^50^, SciPy 1.5^51^ and scikit-image 0.15^52^ for morphological manipulation.

A 3D reconstruction of the t-tubule network was created from confocal NCX1/Cav-3 imaging by first fitting the data to 10 nm x 10 nm x 10 nm voxels, binarizing using an adaptive Gaussian threshold, and then pruning small non-contiguous components (<0.03 μm^3^). The resulting thresholded structures were skeletonized using a 3D skeletonization algorithm^53^, and re-dilated to form 250 nm wide cylindrical t-tubules^54^. The correlated dSTORM imaging data of RyRs were then fitted to the same voxel grid and Otsu-thresholded. The majority of the thresholded clusters directly overlapped with the defined dyadic RyR face. Due to inaccuracies in the t-tubule reconstruction as well as limited precision in the underlying confocal imaging, all clusters lying within 150 nm of a reconstructed t-tubule were estimated to be dyadic clusters. More distally localized RyR clusters were deemed to be non-dyadic.

To render the 3D geometric reconstructions and produce geometric meshes for use in mathematical modeling, specific locations of individual RyRs were assigned by projecting the thresholded dyadic cluster data onto a cylindrical shell surrounding the t-tubules, leaving a 1-voxel wide (10 nm) dyadic cleft. These overlap interfaces were then randomly filled with RyRs such that no channels were closer than a center-to-center distance of 30 nm. As shown in our earlier work, this approach yields RyR numbers that are in good agreement with our experimentally-obtained estimates^5^.

To visualize the reconstructed geometry of t-tubules and RyRs, iso-surfaces were generated from the full geometries using the Lewiner Marching-Cubes algorithm^55^, smoothed using GAMer 2.0^56^, and finally rendered using Blender (Blender Foundation, the Netherlands).

For use in computational modeling, an SR network was heuristically added to the voxel-based reconstruction of t-tubules and RyRs. The junctional SR (jSR) terminals for each cluster were defined by iterative morphological binary dilation of the RyRs constrained to a 20 nm wide cylindrical shell surrounding the t-tubules. Dilation was performed with a structuring element with a square connectivity of one for nine iterations. This method was chosen because it produces a jSR that evenly surrounds the RyR channels in the cluster, defines a well constrained dyadic cleft space, and has a jSR volume on the order of 3-4 × 10^−12^ μl, which is in good agreement with values reported by electron microscopy tomography^57^. The non-junctional network SR (nSR) was added as a regular grid of thin structures throughout the cytosol. While the generated morphology is artificial, the created structure has a volume and surface area in agreement with 3D electron microscopy data^57^ and serves to connect the different CRUs.

For each CRU in the fully reconstructed volumetric geometry, a surrounding region measuring 1 μm x 1 μm x 1 μm was extracted, producing a set of smaller geometries representing individual CRUs for computational modeling of Ca^2+^ release. Roughly one thousand such CRU geometries were extracted and sorted based on the number of contained RyRs, their spatial spread (measured as the root-mean-square of their distance to the cluster center), and the number of individual clusters within the CRU. Four representative geometries were selected containing roughly the same number of RyRs, but with different degrees of compactness. These ranged from a completely solid CRU to a majorly dispersed CRU consisting of numerous, sprawled clusters (Supplementary Table 1).

### Ca^2+^ Imaging and Analysis

Using an LSM 7Live confocal microscope (Zeiss), spontaneous Ca^2+^ sparks were recorded from quiescent cardiomyocytes loaded with fluo-4 AM (20 μmol/L, Molecular Probes, Oregon, USA) and superfused with Hepes-Tyrode solution containing (in mmol/L): 140 NaCl, 0.5 MgCl2, 5.0 HEPES, 5.5 glucose, 0.4 NaH_2_PO_4_, 5.4 KCl and 1.8 CaCl_2_ (pH 7.4, 37°C). Cardiomyocytes were scanned along a 1024 pixel line drawn across the cell’s longitudinal axis, at a temporal resolution of 1.5 ms for a total duration of 6 s. Ca^2+^ sparks were analyzed using a custom program (CaSparks 1.01, D. Ursu, 2003) as previously^25^, with sparks defined as local increases in fluorescence intensity of at least 3 times above background (ΔF/F_0_ ≥ 0.3). Spark characteristics outputted by the program included amplitude (ΔF/F_0_), time to peak (TTP), full duration at half maximum (FDHM), and full width at half maximum (FWHM). Spark frequency was determined by normalizing spark count to the cell length and recording duration.

Ca^2+^ transients were similarly recorded by confocal line-scans in fluo-4 loaded cells, but during field-stimulation via a pair of platinum electrodes (3 ms supra-threshold current pulses at 1 Hz). As previously described^8^, Ca^2+^ release synchrony was calculated by plotting the profile of time to 50% peak fluorescence (TTF_50_) across the cell and measuring the standard deviation of the values. We have termed this measure the ‘dyssynchrony index’. Spontaneous Ca^2+^ waves were measured during pauses in the electrical excitation, and wave frequency was normalized to the recording duration. SR Ca^2+^ content was estimated by measuring the increase in cytosolic Ca^2+^ fluorescence after superfusing cardiomyocytes with 10 mM caffeine (Sigma-Aldrich). SERCA activity was estimated as the difference between the rate constants (1/τ) of the decline of steady-state Ca^2+^ transient (1 Hz) and the caffeine-induced Ca^2+^ transient (1/τ 1Hz − 1/τ caffeine) acquired in the same cell^58^.

### Western Blotting

Frozen tissue from rat left ventricles was homogenized in cold buffer (210 mM sucrose, 2 mM EGTA, 40 mM NaCl, 30 mM HEPES, 5 mM EDTA) with the addition of a Complete EDTA free protease inhibitor cocktail tablet (Roche Diagnostics, Oslo, Norway) and a PhosSTOP tablet (Roche). SDS was then added to the homogenates to a final concentration of 1%, and protein concentrations were quantified using a micro BCA protein assay kit (Thermo Fischer Scientific Inc., Rockford, IL). Bovine serum albumin (BSA) was used as standard protein.

Protein homogenates (5 or 15 μg/lane) were size fractionated on 4–15% or 15% Criterion TGX gels (Biorad Laboratories, Oslo, Norway) and transferred to 0.45 μM PVDF-membranes (GE Healthcare). The membranes were blocked in 5% non-fat milk or 5% Casein (Roche Diagnostics) in Tris-buffered saline with 0.1% Tween (TBS-T) for 1 hr at room temperature, and then incubated with primary antibody overnight at 4°C. The following primary antibodies were employed for immunoblotting: RyR2 (1:1000) (MA3-916, Thermo Fisher Scientific), pSer2808 RyR2 (1:2500) (A010-30, Badrilla), pSer2814 RyR2 (1:2500) (A010-31, Badrilla) and GAPDH (1:500; sc-20357, Santa Cruz Biotechnology). Secondary antibodies were anti-rabbit (NA934V, GE Healthcare) or anti-mouse (NA931V, GE Healthcare). These were incubated for 1 hr at room temperature and blots were developed using Enhanced Chemiluminescence (ECL prime, GE healthcare). Chemiluminescence signals were detected by a LAS 4000 (GE healthcare) and protein levels were quantified using ImageQuant software (GE Healthcare). Results were normalized to GAPDH and then to Sham values.

### Flow cytometry using Fluorescence-activated cell sorting

In order to examine the relative phosphorylation levels of isoproterenol-treated cardiomyocytes, we employed Fluorescence-activated cell sorting. This technique has previously been shown to enable protein quantification in close correlation with results from Western blotting^59^. Fixed cells labelled with primary antibodies against pSer-2808 RyR2 (A010-30, Badrilla) or pSer-2814 RyR2 (A010-31, Badrilla) and secondary AlexaFluor647 antibody were placed in a suspension of PBS with 1% BSA. Flow cytometry analysis was performed on a Sony SH800 Cell Sorter and FlowJo™ Software (Becton, Dickinson and Company; Ashland, OR, USA) was used for analysis. Control treatments were included which contained unlabelled cells, or only secondary antibody. Voltage settings for the side, back, and forward scatter were kept constant for each experiments described. The cell suspension was analysed at a flow rate of 27 μL/min, until >5000 events were captured. A two-step sorting process was employed. Intact cardiomyocytes were separated from other cell types and debris using a dual parameter dot plot for side- and forward scatter (see Supplementary Fig. 2a for representative image). Single cardiomyocytes were separated from doublets using a plot of forward scatter height and pulse width (Supplementary Fig. 2b).

Following sorting, the secondary AlexaFluor647 antibody (A-21237, Thermo Fisher Scientific) in the single cardiomyocyte fraction was excited with the 638 nm laser and emission detected by a photomultiplier tube with a 665/30 Band pass filter. Data are presented as detected fluorescence intensity as a product of cell size. Confirmation of cell viability was performed using LIVE/DEAD™ Fixable Violet Dead Cell Stain (ThermoFisher), excited with the 405 laser and emission detected by via a 450/50 band pass filter.

### Mathematical Modeling of Ca^2+^ Sparks

Effects of changing CRU configuration on Ca^2+^ spark characteristics were interrogated using a mathematical model of Ca^2+^ release in the dyad. This previously described reaction-diffusion model includes independent, stochastic RyR channels coupled with deterministic Ca^2+^ diffusion and buffering^11^. The model was chosen for the current work since it can incorporate changing CRU configurations based on super-resolution imaging. The model was employed as previously described, but with minor adjustments to better align Ca^2+^ release and pump-leak balance behaviour with experimental data from rat and mouse^60-62^. Specifically, we slightly augmented SR Ca^2+^ reuptake by increasing in the amount and density of SERCA, and modestly decreased SR volume and buffering to reduce releasable Ca^2+^. When combined with a slightly more sensitive RyR model^63^, these modifications yielded larger fractional SR Ca^2+^ release, in better agreement with experimental data. Please refer to Supplementary Tables 2 and 3 for a full list of buffering and RyR parameters.

To incorporate the effects of isoproterenol treatment on Ca^2+^ signaling in the CRU, we altered two model parameters; the SR Ca^2+^ load was increased from 900 μM at baseline to 1300 μM, and the RyR opening sensitivity to cytosolic Ca^2+^ was increased by lowering the K_d_ open from 45 μM to 25 μM. These two model parameters and their adjustments were constrained by comparing the amplitude and rise time of simulated sparks to experimental measurements made in control and isoproterenol treatment conditions (Supplementary Fig. 4).

### Analysis of Spark Simulations

400 simulations were performed for both the baseline and isoproterenol parameter sets, with each CRU geometry. The initial conditions were otherwise identical for each simulation. A single, randomly-selected RyR in the CRU was opened, and the simulation was then allowed to progress stochastically. The probability of each RyR opening and closing was thus based on its own locally-sensed Ca^2+^ concentration. The simulations continued until all RyRs in the CRU had been simultaneously closed for at least 1 ms, at which point the simulation was terminated. Thus, the full tail of the spark was not modelled out of consideration for computational efficiency.

To compare the modelled with experimentally-measured sparks, we used the average concentration of Ca^2+^-bound Fluo in the full cytosol of the 1 μm x 1 μm x 1 μm computational domain. This measure was employed instead of more computationally expensive line scan simulations, since our previous work indicated that this simplified approach yields equal predictions of spark amplitude and time course^11^.

For each simulation, we measured the peak amplitude of the Fluo signal and the rise time (TTP). Based on the measured amplitude, we then determined which simulations to consider sparks, using the same threshold as in the experimentally-measured Ca^2+^ sparks (ΔF/F_0_ ≥ 0.3). For simulations where the peak amplitude was below this threshold, we defined sub-spark events as “quarks”^30^ (0.1 < ΔF/F_0_ < 0.3), or “failed sparks” if Ca^2+^ release was negligible (ΔF/F_0_ ≤ 0.1).

Based on the ratio of spark events to sub-spark events, we estimated the probability of a spontaneous opening leading to an observable spark, dubbed the “spark fidelity”. As each individual simulation was an independent stochastic trial in a Bernoulli process, each simulation produced a spark with probability equal to the fidelity. To estimate the fidelity, we used the maximum-likelihood predictor of the probability, which is simply the observed ratio of successful sparks to total simulations. Extra care should be taken in calculating a confidence interval for spark fidelity, as we might expect the fidelity to be very close to or equal to 0 for severely fragmented clusters. We therefore opted to follow the recommendation of Brown, Cai and DasGupta^64^ and used the Agresti-Coull confidence interval^65^.

We also quantified the amount of released Ca^2+^ during each simulation. For simulations that were deemed to elicit observable sparks, the released Ca^2+^ was considered spark-mediated leak, while sub-spark events defined “silent” leak. Total RyR-mediated leak was calculated as the sum of the two types of events. As the mathematical model was mainly constrained using amplitude and TTP values of sparks, the leak measurements were not analyzed in absolute terms, but rather as the relative amount of spark-mediated and silent leak in the different geometry and parameter combinations.

### Statistical Analyses

Cardiomyocytes from rats were randomly selected for analysis of RyR localisation and calcium imaging. Power analysis was performed to determine sample sizes based on known variability of measured parameters. No data were excluded from the analyses, and consistent observations were made during analyses performed on different cardiomyocytes from different hearts. Experiments involving multiple treatment groups were all carried out in the same day to ensure consistency. Statistical significance between sample means was calculated by nested ANOVA (SPSS, IBM), with Fisher’s least significant difference (LSD) post-hoc comparison where appropriate. P values < 0.05 were considered to be significant. All results presented are expressed as mean ± standard error of the mean (SEM) unless otherwise stated.

## Code Availability

Custom code used in this study is available at the public repository https://gitlab.com/louch-group/ryr-tt-correlative-analsyis.

## Acknowledgements

We thank the Section for Comparative Medicine at Oslo University Hospital Ullevål for expert animal care. This study was financially supported by the European Union’s Horizon 2020 research and innovation progamme (Consolidator grant, W.E.L.) under grant agreement No. 647714, the Norwegian Research Council (X.S., W.E.L.), and The Norwegian Association for Public Health (W.E.L.). This work used the Oakforest-PACS supercomputer system provided by The University of Tokyo through Joint Usage/Research Center for Interdisciplinary Large-scale Information Infrastructures and High Performance Computing Infrastructure in Japan (Project IDs: JHPCN-jh180024, JHPCN-jh190040 & JHPCN-jh200036).

## Author Contributions

X.S., A.G.E., C.S. and W.E.L. designed and supervised the study. X.S., J.vdB., A.B-D., A.G.E., C.S. and W.E.L wrote the manuscript. X.S., A.B-D., T.R.K., E.S.N., Y.H., A.P.Q., E.K.S.E. and I.S. performed the experiments. X.S., J.vdB., A.B-D., M.L., E.K.S.E. and I.S. analyzed data. X.S., J.vdB., M.L., I.S., X.H.T.W., A.G.E., C.S. and W.E.L discussed data and gave conceptual advice. All authors critically reviewed the manuscript.

## Abbreviations

β-AR: β-adrenergic receptor
CaMKII: Ca^2+^/calmodulin-dependent kinase II
CICR: Ca^2+^-induced Ca^2+^ release
CRU: Ca^2+^ release unit
dSTORM: Direct Stochastic Optical Reconstruction Microscopy
F_50_: Half-maximal fluorescence
FDHM: Full duration half maximum
FWHM: Full width half maximum
HF: Heart failure
LTCC: L-type Ca^2+^ channel
PKA: Protein kinase A
RyR: Ryanodine Receptor type 2
SR: Sarcoplasmic reticulum
TTF_50_: Time to F_50_
TTP: Time to peak

## SUPPLEMENTARY MATERIAL

### Supplementary Tables

**Supplementary Table 1.** Definition of Modeled RyR Cluster Geometries. From a large number of automatically-generated 3D CRU geometries, four were selected to be used in mathematical modeling of Ca^2+^ release. These were chosen to represent the range of fragmentation observed experimentally, while ensuring that each CRU contained roughly the same number of RyRs. Characteristics for each CRU are listed. Spatial spread was calculated as the root mean square distance of RyRs to the CRU center.

| Geometry Definition | No of RyRs | Spatial Spread | Clusters (No of RyRs / cluster) |
| --- | --- | --- | --- |
| Solid | 35 | 91.2 nm | 1 (35) |
| Minorly Fragmented | 36 | 114.7 nm | 4 (22, 9, 4, 1) |
| Moderately Fragmented | 32 | 135.4 nm | 7 (16, 4, 4, 3, 2, 2, 1) |
| Majorly Fragmented | 34 | 174.4 nm | 5 (12, 10, 5, 4, 3) |

**Supplementary Table 2.** Model parameters for Ca^2+^ buffering.

| | $\sigma$ ($\mu\text{m}^2/\text{s}$) | $B_{\text{tot}}$ | $k_{\text{on}}$ ($\text{ms}^{-1}$) | $k_{\text{off}}$ ($\text{ms}^{-1}$) |
| --- | --- | --- | --- | --- |
| Calmodulin | 22 | 24 $\mu\text{M}$ | 0.034 | 0.238 |
| ATP | 140 | 455 $\mu\text{M}$ | 0.255 | 45 |
| Fluo | 20 | 50 $\mu\text{M}$ | 0.08 | 0.09 |
| Troponin | 0 | 70 $\mu\text{M}$ | 0.0327 | 0.0196 |
| Calsequestrin | 0 | 16 mM | 0.102 | 65 |

**Supplementary Table 3.** RyR model parameters.

| Rates | $k_{\text{min}}$ ($\text{ms}^{-1}$) | $k_{\text{max}}$ ($\text{ms}^{-1}$) | $K_d$ ($\mu\text{M}$) | n |
| --- | --- | --- | --- | --- |
| + | $5 \times 10^{-6}$ | 0.9 | 45 or 25* | 2.7 |
| - | 0.9 | 3 | 45 | -5 |
| Unitary Current | 0.235 pA |  |  |  |

**Supplementary Table 4.** Cardiac parameters from post-myocardial infarction (MI) rats with HF and Sham-operated controls. HF development was examined by CINE MRI assessment, based on established criteria^20^. Infarct size is percentage of left ventricular free wall (Sham: n_heart_ = 14, HF: n_heart_ = 12; *p<0.05).

| Parameter | Sham (n=14) | MI (n=12) |
| --- | --- | --- |
| Infarct size (%) |  | 59.9 ± 2.1 |
| End-diastolic volume (mL) | 0.33 ± 0.01 | 0.03 ± 0.004* |
| End-systolic volume (mL) | 0.09 ± 0.01 | 0.02 ± 0.003* |
| Stroke volume (mL) | 0.24 ± 0.01 | 0.01 ± 0.001* |
| Ejection fraction | 0.73 ± 0.02 | 0.35 ± 0.02* |

### Supplementary Figures

**Supplementary Figure 1:**
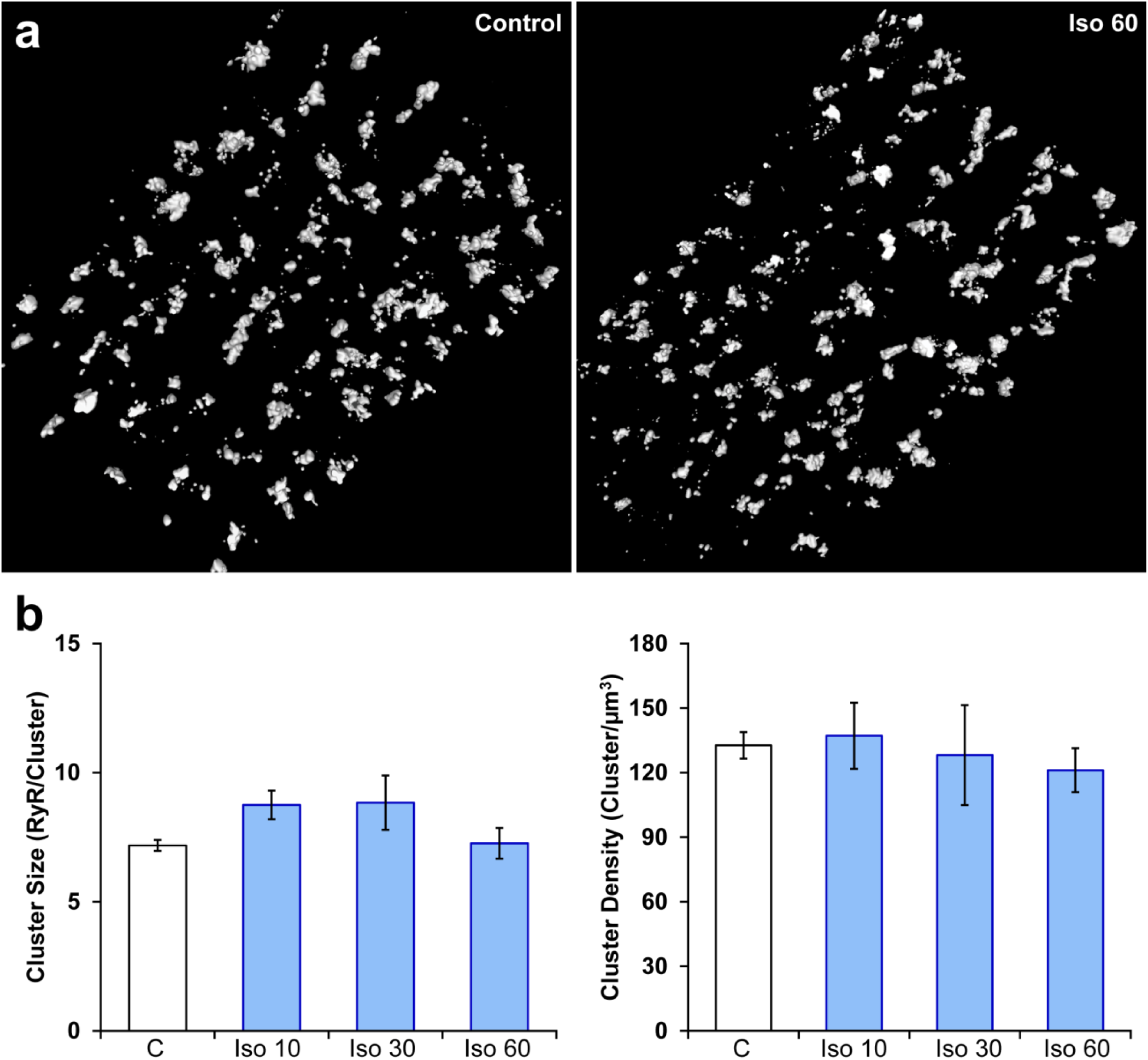
β-AR stimulation does not influence RyR organization on the cell surface. Representative images (**a**) and mean data (**b**) indicate that, in contrast to internal RyRs (**Fig. 1**), RyR clusters remain intact during prolonged β-AR stimulation. (Control: n_cells_ = 32, n_hearts_ = 3; Iso 10: n_cells_ = 14, n_hearts_ = 4; Iso 30: n_cells_ = 12, n_hearts_ = 4; Iso 60: n_cells_ = 24, n_hearts_ = 4).

**Supplementary Figure 2:**
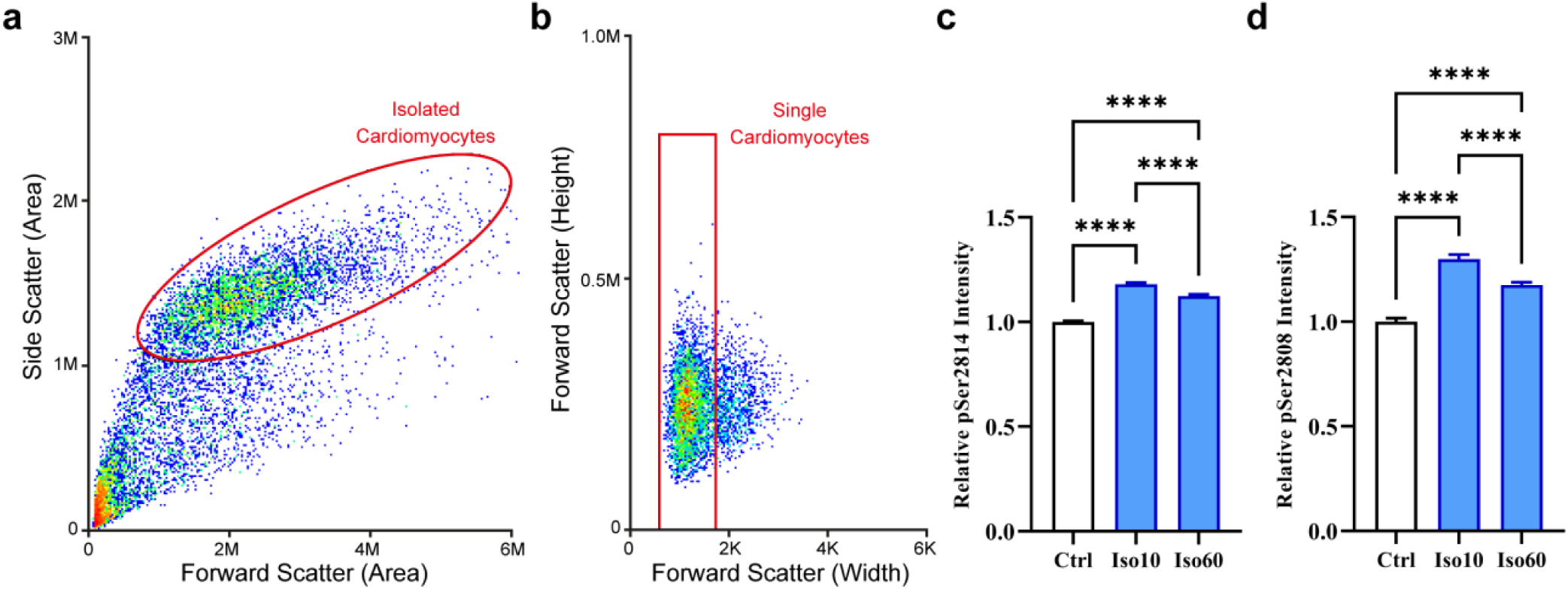
Quantification of RyR phosphorylation by flow cytometry. Cardiomyocytes were separated from other cell types and debris using a dual parameter dot plot for side and forward scatter (**a**), and the single cardiomyocyte fraction was identified using a plot of forward scatter height and pulse width (**b**). (**c**) Fluorescent detection of RyR phosphorylation at ser-2814 revealed increased phosphorylation during 1 h isoproterenol treatment period, which was most prominent at the early time point examined (Control: n_cells_ = 7140, Iso 10: n_cells_ = 6251,; Iso 60: n_cells_ = 5253; n_hearts_ = 2). (**d**) RyR phosphorylation at ser-2808 also exhibited a bell-shaped response to isoproterenol treatment, but remained increased after 1 h of treatment. (Control: n_cells_ = 742, Iso 10: n_cells_ = 803,; Iso 60: n_cells_ = 1630; n_hearts_ = 1).

**Supplementary Figure 3:**
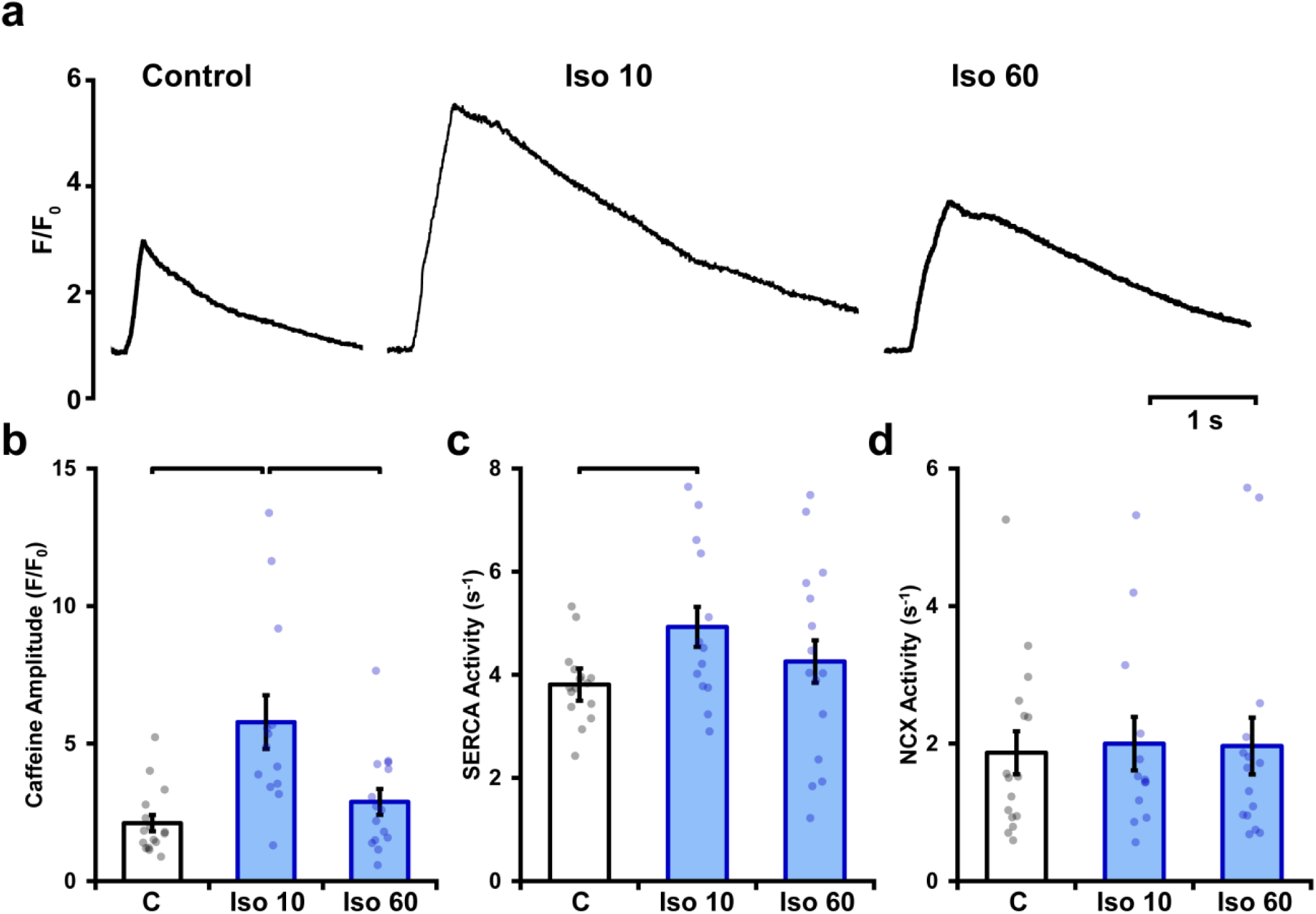
Isoproterenol elicits a biphasic increase in SR Ca^2+^ content. SR Ca^**2+**^ content was estimated by rapidly changing the superfusate to one containing 10 mmol/L caffeine, and measuring the amplitude of the elicited Ca^**2+**^ release. Representative recordings (**a**) and mean data (**b**) show that a marked increase in SR content during early isoproterenol treatment was reversed with continued exposure. SERCA and NCX activity (**c** and **d**, respectively) were estimated based on fits of the decays of caffeine- and electrically-elicted Ca^**2+**^ transients (see methods). (Control: n_cells_ = 16, n_hearts_ = 4; Iso 10: n_cells_ = 13, n_hearts_ = 4; Iso 60: n_cells_ = 15, n_hearts_ = 4).

**Supplementary Figure 4:**
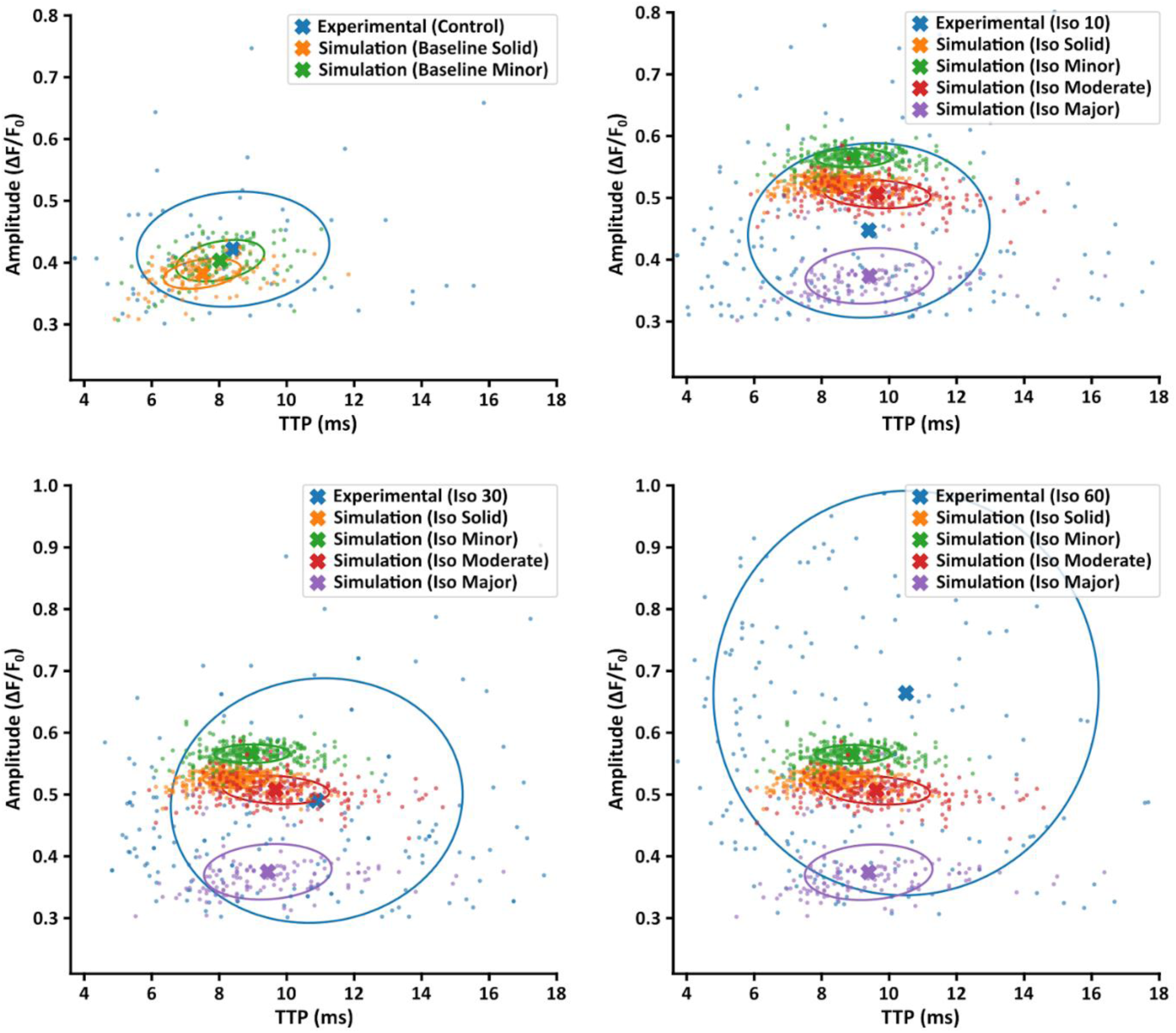
Comparison of experimentally-measured and simulated Ca^2+^ sparks. Amplitude and time to peak (TTP) values for experimentally measured Ca^2+^ sparks are plotted under control conditions and the three time points of isoproterenol stimulation (10, 30, 60 min). Comparison is made with simulated sparks generated by the four modelled CRU geometries under baseline conditions, and under simulated isoproterenol conditions including increased RyR release flux and Ca^2+^ sensitivity. Note that the simulated sparks shown are identical in the three latter plots. Crosses indicate population means and circles show the covariance ellipse at 1 standard deviation.

**Supplementary Figure 5:**
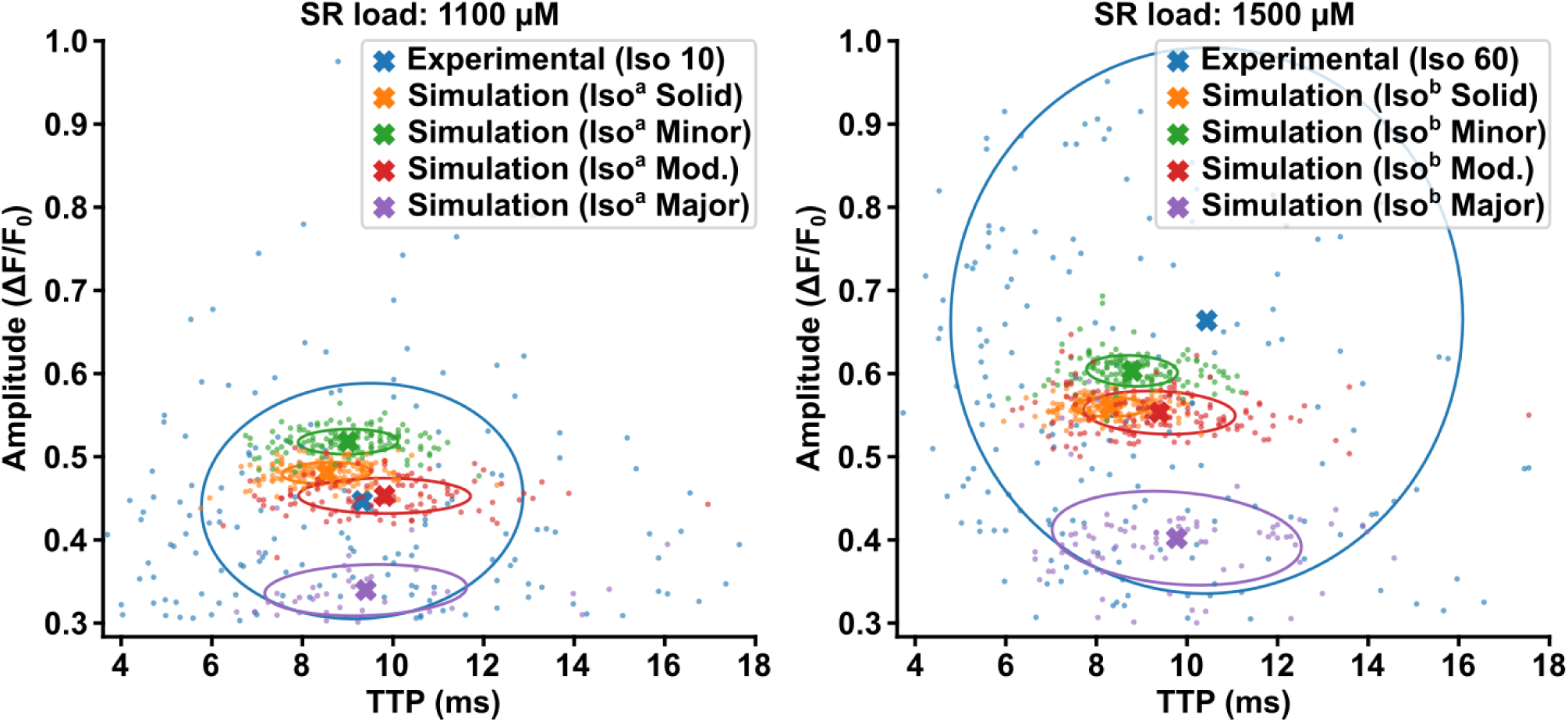
Model behavior at different SR loads. To test model behavior, we perturbed the SR load parameter in the isoproterenol model while maintaining RyR Ca^2+^ sensitivity. We compared the standard isoproterenol model (ie. SR load = 1300 μM, see Supplementary Fig. 2) with SR content reduced to 1100 μM (left panel, “Iso^a^”), or increased to 1500 μM (right panel, “Iso^b^”). These perturbations had little effect on spark TTP, with similar trends observed across the modeled CRU geometries. Spark amplitude, on the other hand, was observed to be markedly sensitive to SR load perturbation. Results are plotted alongside experimental data recorded at 10 min or 60 min following isoproterenol treatment (left and right panels, respectively).

**Supplementary Figure 6.**
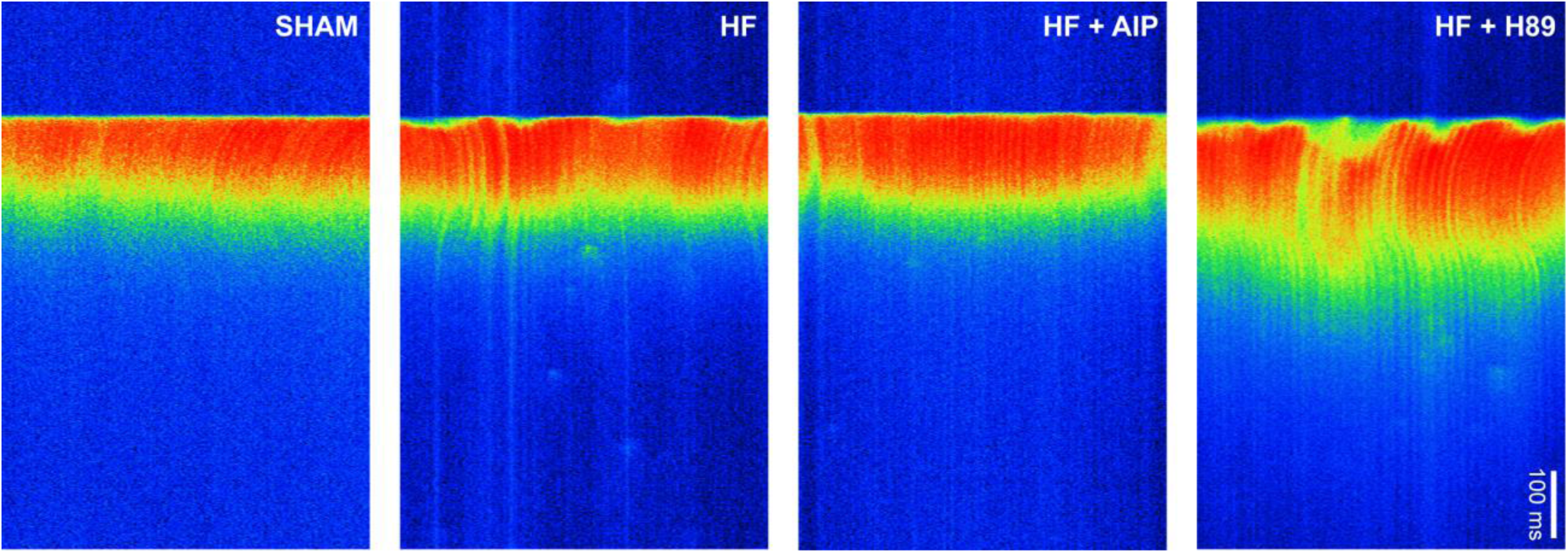
Ca^2+^ transients in Sham and HF myocytes. Representative linescan images illustrating the last in a series of electrically-paced Ca^2+^ transients. Dyssynchronous and slowed Ca^2+^ release observed in HF myocytes was reversed by treatment with AIP but not with H89. See Fig. 6m-o for mean measurements.

